# *Trichoderma asperelloides* enhances local and systemic acquired resistance response under low nitrate nutrition in Arabidopsis

**DOI:** 10.1101/502492

**Authors:** Aakanksha Wany, Pradeep K. Pathak, Alisdair R Fernie, Kapuganti Jagadis Gupta

## Abstract

Nitrogen (N) is essential for growth, development and defense but, how low N affects defense and the role of *Trichoderma* in enhancing defense under low nitrate is not known. Low nitrate fed Arabidopsis plants displayed reduced growth and compromised local and systemic acquired resistance responses when infected with both avirulent and virulent *Pseudomonas syringae* DC3000. These responses were enhanced in the presence of *Trichoderma*. The mechanism of increased local and systemic acquired resistance mediated by *Trichoderma* involved increased N uptake and enhanced protein levels via modulation of nitrate transporter genes. The *nrt2.1* mutant is compromised in local and systemic acquired resistance responses suggesting a link between enhanced N transport and defense. Enhanced N uptake was mediated by *Trichoderma* elicited nitric oxide (NO). Low NO producing *nia1,2* mutant and *nsHb^+^* over expressing lines were unable to induce nitrate transporters and thereby compromised defense in the presence of *Trichoderma* under low N suggesting a signaling role of *Trichoderma* elicited NO. *Trichoderma* also induced SA and defense gene expression under low N. The SA deficient *NahG* transgenic line and the *npr1* mutant were also compromised in *Trichoderma*-mediated local and systemic acquired resistance responses. Collectively our results indicated that the mechanism of enhanced plant defense under low N mediated by *Trichoderma* involves NO, ROS, SA production as well as the induction of NRT and marker genes for systemic acquired resistance.

**One-sentence summary:** *Trichoderma* enhances local and systemic acquired resistance under low nitrate nutrition

## Introduction

Nitrogen (N) is essential for growth and development of plants. It is a crucial component in chlorophyll, nucleic acids and amino acids as well as a large number of specialized metabolites. N, furthermore, plays an important role in the regulation of primary and secondary metabolism and also in protection of plants against biotic and abiotic stresses since it triggers defense response against stresses (O’Brien *et al*. 2016; Mur *et al*. 2017). N deficiency occurs in soil due to slow mineralization, lack of sufficient organic matter, leaching due to heavy rainfalls and increased activities of denitrifying bacteria. Low N can retard plant growth and cause severe physiological and morphological defects (Walker *et al*. 2001; Landrein *et al*. 2018). To cope with this, plants have evolved N uptake systems that support their survival under N deficiency (Li *et al*. 2017). These uptake systems are based on their affinity with transport of nitrates (NO_3_^−^), being mediated by the low and high affinity nitrate transporter protein families (LATs & HATs; Tsay *et al*. 2007).

N plays a very significant role in plant defense against both virulent and avirulent pathogens (Dietrich *et al*. 2004; Gupta *et al*. 2013; Mur *et al*. 2017). Hence, operation of an efficient N transport system can help plants to obtain sufficient nitrogen to defend themselves against bacterial pathogens. These defense responses involve activation of innate immune response comprising of pathogen-triggered immunity (PTI and effector-triggered immunity (ETI) (Alves *et al*. 2014; Jones and Dangl 2006) and require N. One of the defense mechanisms against invading pathogen importantly includes rapid programmed cell death known as the hypersensitive response (HR), which develops during incompatible plant-pathogen (*R-avr*) interactions (Delledonne *et al*. 1998). An early characteristic of HR is the rapid generation of superoxide (O_2_^−.^), nitric oxide (NO) and the accumulation of H_2_O_2_ (Lamb and Dixon, 1997). Moreover, SA produced during HR, plays an important role in plant defense (Mur *et al*. 2000, 2008; Gupta *et al*. 2013). The host mobilizes salicylic acid (SA) for plant defense (Oliva and Quibod 2017, Mur *et al*. 2017) and this pathway itself requires N for synthesis of various intermediates leading to SA production. Thus, N not only improves the nutritional status of the plant, but, also it also plays a more direct role in defense. Moreover, the form of N nutrition can greatly influence the HR-mediated resistance in plants (Gupta *et al*. 2013).

Nitrate (NO_3_^−^) nutrition greatly influences HR via the production of NO which is a regulatory signal in plant defense (Delledonne *et al*. 1998). NO production depends on NO_3_^−^ or L-Arginine (Planchet *et al*. 2005, Astier *et al*. 2018) As such, N deficiency can also lead to reduced levels of NO. Moreover, SA is known to be induced by NO, hence, low NO_3_^−^ leads both to low NO and reduced SA levels. Since NO production also requires NO_3_^−^, it may also play a role in local acquired resistance (LAR) where resistance is developed at the site of infection and also in systemic acquired resistance (SAR) which results in broad-spectrum disease resistance against secondary infections following the primary infection (Cameron *et al*. 1994, 1999). SAR develops either as a consequence of HR where NO has a proven role (Delledonne *et al*. 1998). SAR is dependent on SA or its derivatives (Park *et al*. 2007, Metraux *et al*. 1990) as well as pathogen responsive (*PR*) gene expression (Ryals *et al*. 1996). It was previously shown that NO plays a role in nitrate uptake/assimilation during stress by modulation of the expression of nitrate transporters (Frungillo *et al*. 2013). Thus, N plays likely plays multifaceted roles in plant defense. Indeed, whilst it has been shown that a very high levels of N can increase susceptibility to pathogen infection (Fagard *et al*. 2014) a low level of N can also increase susceptibility (Dietrich *et al*. 2004). For this reason we set out to test whether increasing N uptake can assist in plant defense under low NO_3_^−^. Many plant symbiotic microbes such as mycorrhiza, *Trichoderma* and plant growth promoting rhizobia are known to increase nutrient uptake. However, the operation of plant defense under low N and the effect of these microbes in increasing plant defense under N deficiency is currently unknown.

Several species of *Trichoderma* play an important role in plant growth promotion and resistance against various biotic and abiotic stresses. They confer resistance to plants via various mechanisms such as mycoparasitism, activation of basal and induced systemic resistance (ISR) responses (Brotman *et al*. 2012). Upon pathogen attack, *Trichoderma* treated plants show both elevated elevated phenylalanine ammonia-lyase (*PAL*) transcript levels (Yedidia *et al*. 2003) and increased levels of defense-related plant enzymes (Shoresh *et al*. 2005).

If *Trichoderma* enhances N uptake, it could be anticipated to have positive effects on plant defense via both LAR and SAR responses. So far is it not known whether *Trichoderma* can increase plant defense under low N. Therefore, in the present study, we have assessed, the systemic defense response of low N fed *Arabidopsis thaliana* plants to the phytopathogen; *Pseudomonas syringae* p.v. tomato DC3000 induced by the beneficial fungus *Trichoderma asperelloides* (*T203*) under low and optimum nitrate concentrations. Here, we describe that *Trichoderma* enhances the SAR response in Arabidopsis grown under low N by enhancing NO_3_^−^ uptake via enhancing nitrate transporter expression with this mechanism involving both NO and SA.

## Results

### Low nitrogen compromises LAR response and *Trichoderma* enhances LAR under low N condition

Nitrogen plays an important role in plant defense, hence, we first tested the effect of low nitrate on the LAR response. For this purpose, WT plants were grown under high (3 mM) and low (0.1 mM) NO_3_^−^ conditions and infected with avirulent *P. syringae* DC3000 (*avrRpm1*) and the local HR response was observed (Fig. **1a**, Fig. **S3b**). As shown in Fig.**1a-WT panel**, the 3 mM NO_3_^−^-fed *Pst* infiltrated plants showed HR at the inoculation sites at 24 and 48 hours post infection (hpi) whereas plants grown in the presence of 0.1 mM NO_3_^−^ plants displayed chlorotic lesions at the site of inoculation at 24 and 48 hpi. We next studied the effect of *Trichoderma* on the LAR response Surprisingly, 0.1 mM NO_3_^−^ grown WT plants in the presence of *Trichoderma* did not show any severe disease symptoms, rather they mimicked the HR phenotype of 3 mM NO_3_^−^ grown plants and displayed resistance (Fig. **1a-WT+T panel**). Interestingly, we also found that under 0.1 mM NO_3_^−^, *Trichoderma* treated plants displayed an enhanced growth phenotype compared to plants in the absence of *Trichoderma* (Fig. **S1a**), this enhanced growth was linked to an increased leaf number, leaf fresh weight and total chlorophyll content (Fig. **S2 b, c, d**). These results suggest that *Trichoderma* provides resistance under low N stress. Furthermore, HR was evidenced in electrolyte leakage (EL) assays where higher and more rapid leakage was recorded in the presence of *Trichoderma* irrespective of the concentration of nitrate supplied (Fig.**1b**) with these results being in close accordance with the observed HR response.

**Fig 1:**
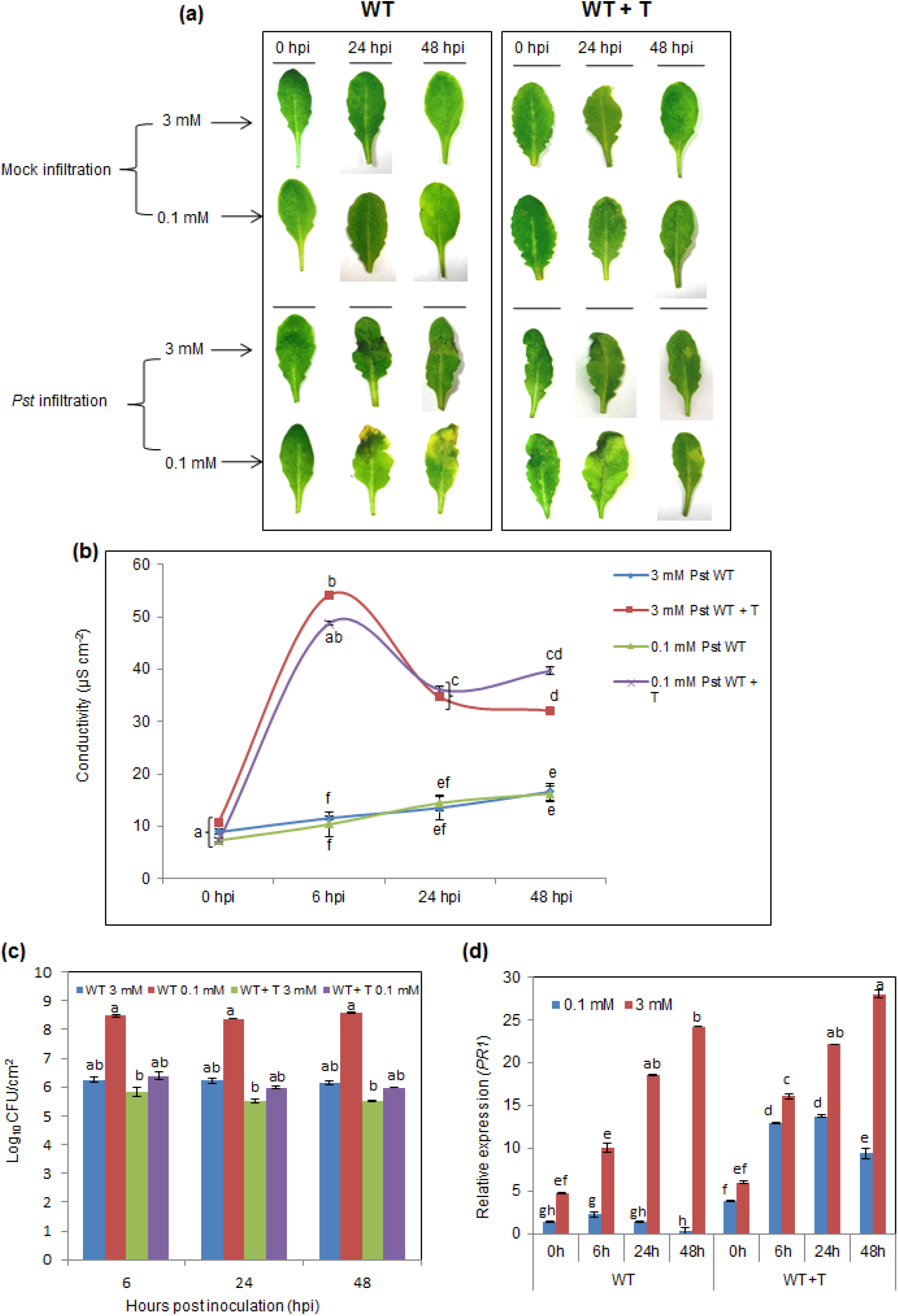
*Trichoderma* supplementation enhances LAR response elicited by *Pst*DC3000/avrRpm1 in low nitrate grown WT plants. LAR response shown by WT and WT+T grown plants fed with 0.1 mM and 3 mM nitrate nutrition subjected to *Pst*DC3000/avrRpm1 infection. Two sets of plants viz., control WT plants for *Pst* treatment were grown in autoclaved un-inoculated soilrite: agropeat mixture and treated WT plants were grown in pots with soilrite: agropeat with *Trichoderma* spore suspension and denoted as WT+T plants. (a) *Pst*DC3000-avrRpm1 mediated HR phenotype observed in 3 mM and 0.1 mM NO_3_^−^ grown WT and WT+ T plants at different time points. First panel shows the mock infiltrated leaves i.e., control leaves treated with 10 mM MgCl_2_ and second panel shows *Pst* infiltrated leaves. Image is representative of 4 independent replicates. (b) Electrolyte leakage from infiltrated leaf areas with *Pst* in 0.1 mM and 3 mM NO_3_^−^ grown WT and WT + T plants at different time points. Line graph represents the average of five biological replicates ± SEM. (c) Bacterial number in log CFU in *Pst* infiltrated leaves of WT and WT+T plants at different time points. Column graph represents the average of three biological replicates ± SEM. (d) Relative *PR1* transcript levels from WT and WT+T plants at different time points. Data are average mean values ± SE with n=3. In all the experiments, statistical significance was tested by two-way ANOVA followed by Tukey’s All-Pairwise Comparisons post-hoc test. The different letters above each column represent significance difference between means at p□<□0.05.

A significant increase in the colony forming unit (CFU) count during LAR was observed in 0.1 mM WT plants supplied with 0.1 mM nitrate was observed in comparison to the bacterial numbers of the same plants grown in the presence of *Trichoderma* (Fig. **1c**). We further found that, *PR1* (a marker for HR) transcript levels increased in *Pst* treated plants grown in the presence of either 0.1 mM (~9 fold at 48 hpi) or 3 mM (~28 folds at 48 hpi) NO_3_^−^ in the presence of *Trichoderma* grown in comparison to WT plants grown on 0.1 mM NO_3_^−^ and in the absence of *Trichoderma*, where expression of this gene remained unaltered at 48 hpi (Fig.**1d**). Collectively these data thus suggest that *Trichoderma asperelloides* plays a role in improving the LAR response under low NO_3_^−^ availability.

### *Trichoderma* can induce the SAR response under low N

Having characterized LAR responses, we next studied responses SAR by infecting the secondary leaves with virulent *Pst* DC3000 (Fig. **S1b**; Cameron *et al*. 1994 and 1999). Three days post-secondary challenge with virulent *Pst* DC3000, EL, CFU and *PR1* gene expression were assessed (Fig. S3c). During this SAR assay *Pst*-challenged WT plants grown on 0.1 mM NO_3_ showed more prominent necrotic and dark yellow lesions that spread to both sides of the leaf than *Pst*-challenged WT plants grown on 3 mM NO_3_. This suggests that, the plants grown on 3 mM NO_3_^−^ had stronger defenses during secondary challenge than the plants grown on 0.1 mM NO_3_^−^ suggesting that sufficient N is required for SAR development. In the presence of *Trichoderma* there was significant reduction of yellowing in the 0.1 mM NO_3_^−^ plants in comparison to plants in which *Trichoderma* was absent (Compare Fig. **2a-WT+T panel** with **WT panel**).

**Fig. 2.**
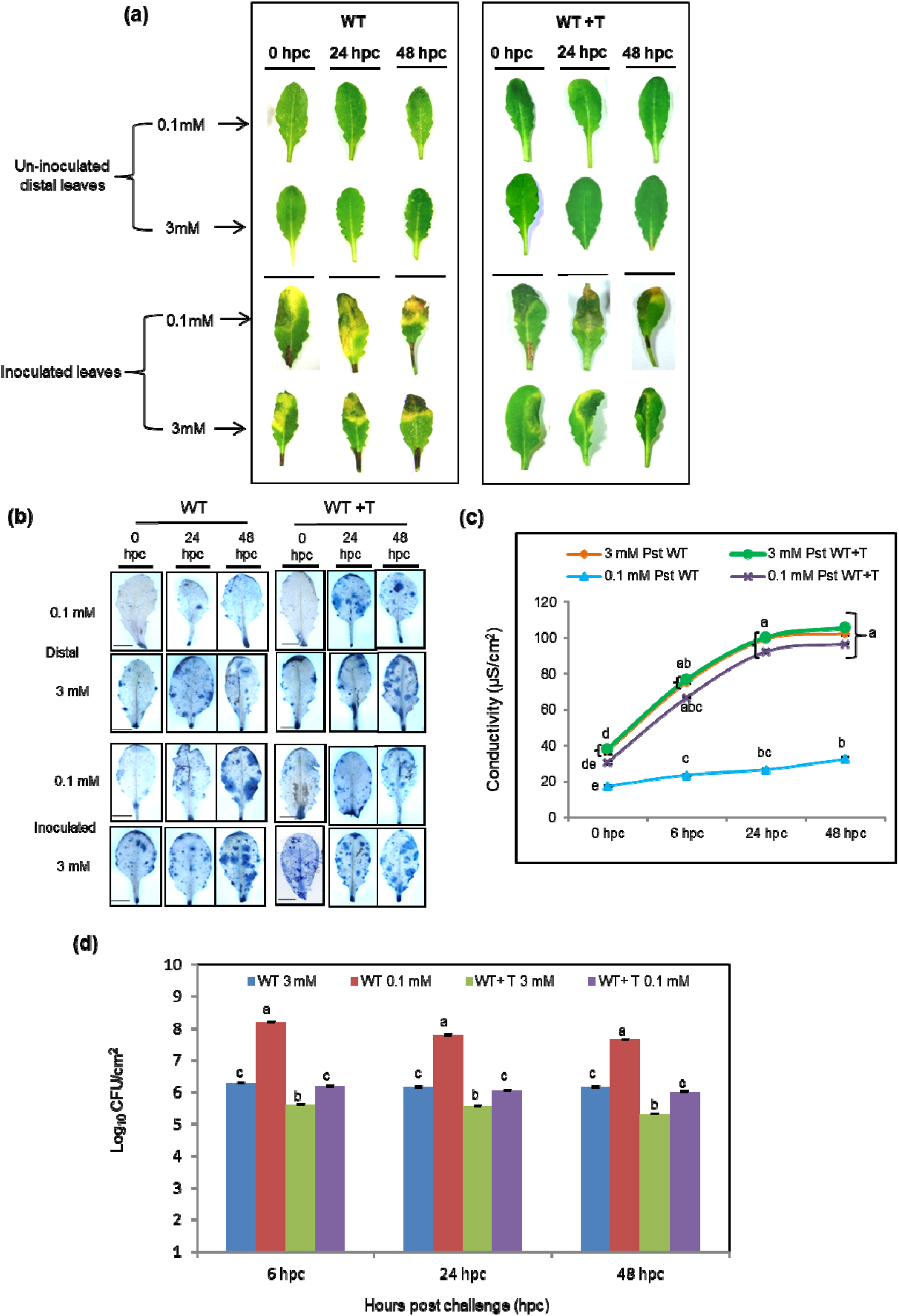
*Trichoderma* enhances SAR response in low nitrate grown WT plants. SAR response shown by WT and WT+T plants grown in 0.1 mM and 3 mM nitrate nutrition were initially subjected to primary inoculation on one leaf per plant with *Pst*DC3000/avrRpm1, left for two days followed by secondary challenge inoculation with virulent *P. syringae* on 4 other (distal) leaves per plant, leaving 5-6 healthy un-inoculated leaves per plant (Cameron et al. 1999). These plants are left for 3 days and then the Inoculated (I) and un-inoculated/distal (U) leaves were assayed for HR phenotype, Trypan blue imaging, EL pattern and bacterial count experiments. The uninoculated leaves from now is mentioned as distal leaves. (a) HR phenotype post-secondary challenge in inoculated and uninoculated leaves of WT and WT+T plants at different time points in 0.1 mM and 3 mM NO_3_^−^. (b) Histochemical staining using Trypan Blue for the detection of HR-mediated cell death post-secondary challenge in inoculated leaves of WT and WT+T plants at different time points in 0.1 mM and 3 mM NO_3_^−^. Images were observed under the 0.5 X objective of AZ100 Stereo Microscope. Scale bar-1 mm. (c) Electrolyte leakage observed post-secondary challenge in inoculated and uninoculated leaves of WT and WT+T plants at different time points in 0.1 mM and 3 mM NO_3_^−^. (d) Bacterial number in log CFU post-secondary challenge in inoculated leaves of WT and WT+T plants at different time points in 0.1 mM and 3 mM NO_3_^−^. Data are average mean values ± SE with n=3. In all the experiments, statistical significance was tested by two-way ANOVA followed by Tukey’s All-Pairwise Comparisons post-hoc test. The different letters above each column represent significance difference between means at p□<□0.05.

Trypan blue images of inoculated and distal leaves are shown in Fig. **2b** and **Fig. S5b**. In response to *Trichoderma*, plants grown on 0.1 mM NO_3_^−^ showed more trypan blue spots during secondary challenge. Upon infection, cell death spots were more widely spread in these leaves (compare Fig. **2b WT+T panel** with **WT panel**). At 48 hpc, there was an increased cell death in inoculated WT leaves grown both on 0.1 and 3 mM NO_3_^−^ but the cell death phenotype differs. The distal leaves plants in the presence of *Trichoderma*, showed a uniform spread of microbursts throughout the leaf blades, at all the time points but with the plants grown on 3 mM NO_3_^−^ displaying slightly larger microbursts than those grown on 0.1 mM NO_3_^−^ (Fig. **2b**). The presence of these microbursts suggests that during SAR establishment the mobile signal perceived by the distal leaves results in the occurrence of low frequency microscopic HRs (Cameron *et al*. 1999; Alvarez *et al*. 1998).

We next investigated electrolyte leakage during SAR. Mock inoculated plants showed moderate EL in plants grown on either 0.1 mM or 3 mM NO_3_^−^ (Fig. **S2a**). EL was, however, significantly higher (~4 fold) in both 0.1 and 3 mM NO_3_^−^ grown plants treated with *Trichoderma* during secondary challenge, in comparison to 0.1 mM nitrate grown plants in the absence of *Trichoderma* at 24 and 48 hpc (Fig. **2c**).This suggests that *Trichoderma* treatment primes the defense responses in the leaves. EL data of distal leaves during SAR is also shown in Fig. **S5a**.

Bacterial count in terms of Log CFU (Fig. **2d**) revealed that, the *Pst* population significantly decreased in *Trichoderma* treated low N-fed plants, suggesting that *Trichoderma* imparts enhanced systemic resistance to the NO_3_^−^ stressed plants.

### *Trichoderma* activates nitrate transporters, facilitates N uptake and promotes SAR in low N-fed plants

Increased LAR and SAR responses under low NO_3_^−^ in the presence of *Trichoderma* are probably due to an increased nutrient uptake as these responses were all absent in the absence of *Trichoderma*. We, therefore, checked the NO_3_^−^ levels in the leaves of 0.1 mM and 3 mM NO_3_^−^ grown WT plants in the presence and absence of *Trichoderma* and represented it as percent nitrate uptake up to Day 15 in leaves (Fig. 3a). The experimental design and protocol for measuring NO_3_^−^ levels and uptake in leaves is demonstrated in Fig. **S4a, b** and **d**. It was observed that % nitrate uptake was significantly increased in the presence of *Trichoderma* in comparison to in the absence of *Trichoderma* enhancing the nitrate uptake of 0.1 mM NO_3_^−^ fed WT plants by 17%. The basal levels of NO_3_^−^ present in the soilrite: agropeat mixture is also shown in Fig **S4c**. This increased NO_3_^−^ transport is probably the reason for resistance hence, we next checked the expression of both LATs chloride channel *CLC-A*), *NRT1/PTR* family 1.2 (*NPF1.2*) and high affinity nitrate transporters (HATs; *NRT2.1, NRT2.2* and *NRT2.4*).

**Fig. 3:**
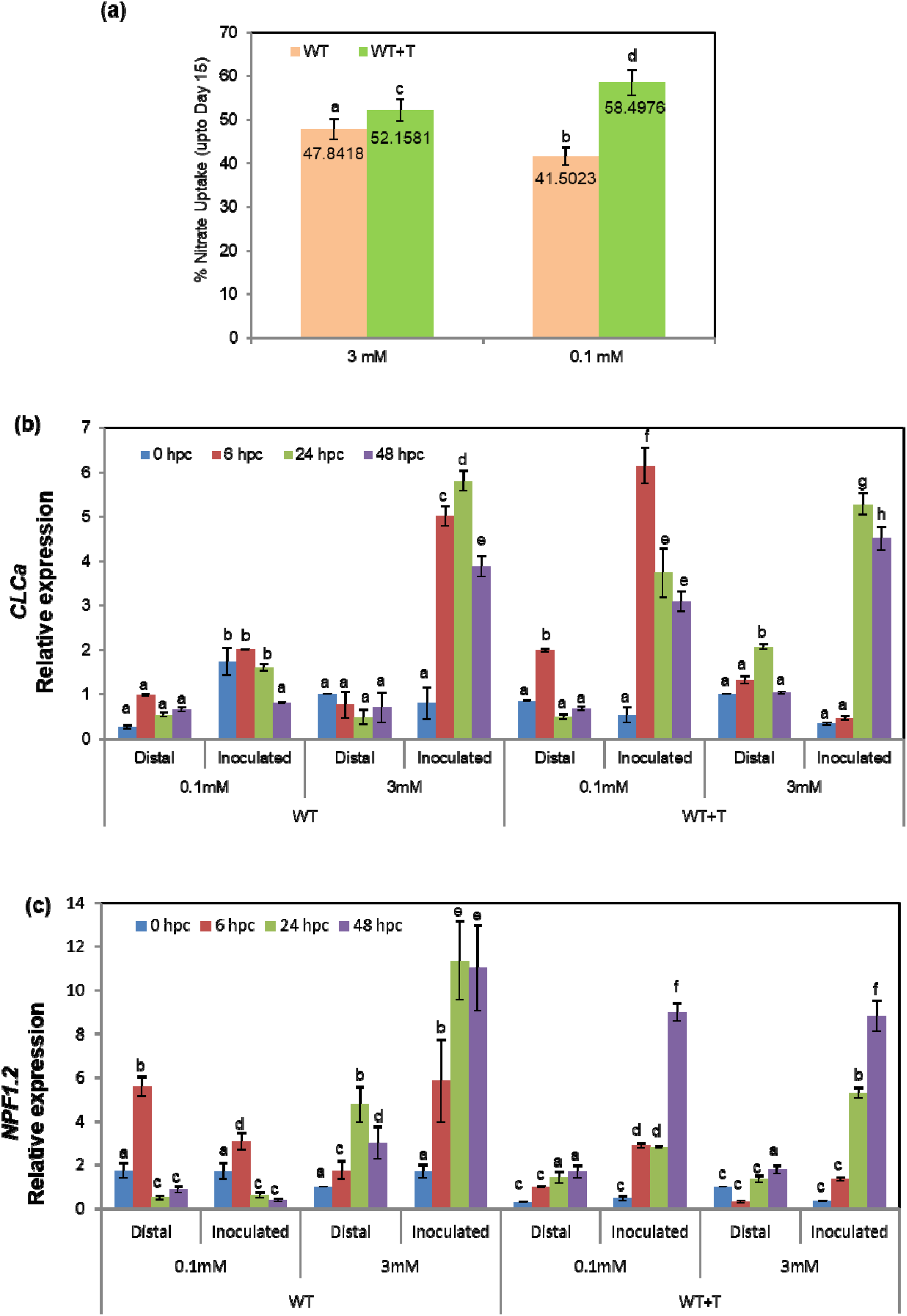
Nitrate uptake and expression analysis of low affinity nitrate transporters. **(a)** Column graph represents the % nitrate levels utilized by the plants till 15th day from the initial day in 0.1 mM and 3 mM WT and WT+T leaves. Data are average mean values ± SE with n=3. Statistical significance was tested by two-way ANOVA followed by Tukey’s All-Pairwise Comparisons post-hoc test. The different letters above each column represent significance difference between means at p□<□0.05. **Expression profile of low affinity nitrate transporter genes (LATs)** during SAR. Relative expression of **(b)** *CLCA* gene in WT and WT+T plants and **(c)** *NPF1.2* gene in WT and WT+T plants grown under 0.1 mM and 3 mM NO_3_^−^ concentration post secondary challenge in both inoculated and distal leaves. For all the target genes, fold expression values are means (n=3) ± SE. Statistical significance was tested by two-way ANOVA followed by Tukey’s All-Pairwise Comparisons post-hoc test. The different letters above each column represent significance difference between means at p□<□0.05.

*CLCA* is a tonoplast located antiporter channel system which drives NO_3_^−^ accumulation in the vacuoles (Krapp *et al*. 2014). It was observed that, under low NO_3_^−^, *CLCA* is less inducible in *Pst* treated plants whereas it was significantly upregulated in 0.1 mM NO_3_^−^ grown, *Pst* treated plants in the presence of *Trichoderma* (Fig. 3b). We further checked the expression of *NPF1.2;* (Fig. **3c**) which is involved in the transfer of xylem-borne nitrate to the phloem in the petiole (Krapp *et al*. 2014). It was found that, in *Pst* treated 3 mM NO_3_^−^ grown WT plants, *NPF1.2* levels were highly induced till 48 hpc, in comparison to 0.1 mM *Pst* inoculated leaves (Fig. **3c**). This might be the reason behind the resistance of 3 mM plants and susceptibility of 0.1 mM plants following pathogen challenge. However, upon *Trichoderma* pre-treatment, *NPF1.2* transcript levels in *Pst*-inoculated 0.1 mM NO_3_^−^ grown plants, showed significant induction in comparison to same treatment in the absence of *Trichoderma*. This suggests that, *Trichoderma* may cause the regulation of *NPF1.2*.

*NRT2* transporters (*NRT2.1, NRT2.2* and *NRT2.4*) are HATS which become activated at low N concentrations (<1 mM). Previously, it was shown that *NRT2.1* is active only under conditions of N starvation (Dechorgnat *et al*. 2012). Under low NO_3_^−^ conditions, *NRT2.1* was significantly induced in all treatments in the presence of *Trichoderma* in *Pst*-treated plants in comparison to plant in the absence of *Trichoderma* where there was already an induction of this gene at early time points (Fig. **4a**). The SAR establishment stage represented by non-inoculated leaves (distal) showed dynamic and elevated up-regulation of *NRT2.1* transcripts (Fig. **4a**). This revealed that, *Trichoderma* colonization benefits the plant by facilitating the N supply even under low NO_3_^−^ conditions thereby improving plant defense via the mediation of SAR. *NRT2.2* expression levels have increased several folds in 0.1 mM distal and inoculated leaves of *Trichoderma* grown plants (Fig. **4b** **& 6b)**). This again suggests that faster N uptake by *Trichoderma* treated plants grown on low NO_3_^−^ can aid in *Pst* defense. We next checked *NRT2.4* expression levels across the treatments. This gene showed early induction (6 h) in 0.1 mM *Pst* treated in inoculated and distal leaves (Fig. **4c**) and the early induction was similar in *Trichoderma* treatment.

**Fig. 4:**
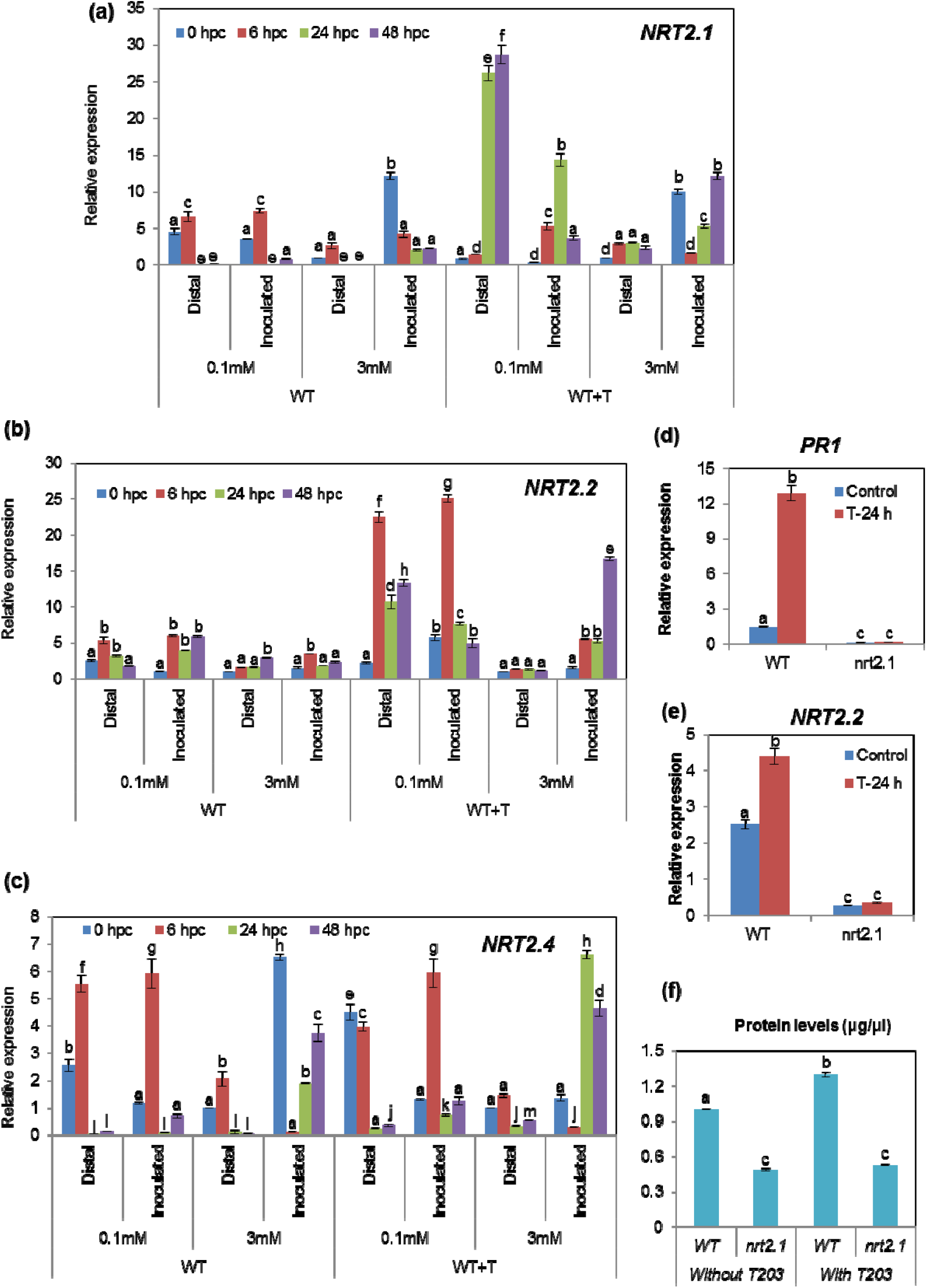
Expression profile of high affinity nitrate transporter genes (HATs) during SAR and response of genotypes (WT and *nrt2.1*) on priming effect of 24 h *Trichoderma* pre-treatment w.r.t *PR1* and *NRT2.2* expression on low nitrate fed WT plants. Relative expression of (a) *NRT2.1*, (b) *NRT2.2*, (c) *NRT2.4* in WT and WT+T plants grown under 0.1 mM and 3 mM NO_3_^−^ concentration post-secondary challenge in both inoculated and distal leaves. For all the target genes, fold expression values are means (n=3) ± SE. Statistical significance was tested by two-way ANOVA followed by Tukey’s All-Pairwise Comparisons post-hoc test. The different letters above each column represent significance difference between means at p□<□0.05. (d) Relative *PR1* expression in roots of WT and *nrt2.1* plants grown under 0.1 mM NO_3_^−^ concentration (with and without *Trichoderma* treatment; 24 h). Data are average mean values ± SE with n=3. Statistical significance was tested by one-way ANOVA followed by Dunnett’s multiple comparison test. The different letters above each column represent significance difference between means at p□<□0.05. (e) Relative *NRT2.2* expression in roots of WT and *nrt2.1* plants grown under 0.1 mM NO_3_^−^ concentration (with and without *Trichoderma* treatment; 24h). Data are average mean values ± SE with n=3. Statistical significance was tested by one-way ANOVA followed by Dunnett’s multiple comparison test. The different letters above each column represent significance difference between means at p□<□0.05. (f) Protein levels in WT and *nrt2.1* plants grown under 0.1 mM NO_3_^−^ concentration in the presence or absence of *Trichoderma*. Data are average mean values ± SE with n=3. Statistical significance was tested by one-way ANOVA followed by Dunnett’s multiple comparison test. The different letters above each column represent significance difference between means at p□<□0.05.

Taken together, these data suggest that *Trichoderma* induces HATs (*NRT 2.1 and NRT 2.2*) to facilitate N uptake under low NO_3_^−^ conditions. To confirm the role of HATs in increasing plant defense via N uptake, we checked the *PR1* expression in WT and *nrt2.1* mutants. The expression of *PR1* gene in *Trichoderma* inoculated plants increased 12-13 folds in 24 h but in the case of *nrt2.1* mutant (Fig. **4d**), it was not at all induced suggesting that *NRT2.1* plays an important role in increasing plant defense under low N mediated by *Trichoderma*. We subsequently found that the expression of the *NRT2.2* gene was also suppressed in the *nrt2.1* mutant, both in and the presence and absence of *Trichoderma* (Fig. **4e**) suggesting that *NRT2.1* is mainly responsible for increasing N uptake under low NO_3_^−^ facilitated by *Trichoderma*. We next checked the protein levels in WT and the *nrt2.1* mutant in the presence or absence of *Trichoderma* under low NO_3_^−^ nutrition. In response to *Trichoderma*, protein levels were increased in the WT. In the *nrt2.1* mutant protein levels were lower than in WT and upon *Trichoderma* treatment only a slight increase in their levels was observed (Fig. **4f**). These results suggest that *Trichoderma* application can increase NO_3_^−^ transport.

### *Trichoderma* elicits NO production during early stages of inoculation which is required for induction of HATs and *PR* gene expression

Previously, it was shown that *Trichoderma* elicits NO at early stages (Gupta *et al*. 2014). This elicitation could subsequently play a role in the induction of NO_3_^−^ transporter genes. Hence, we checked NO production in WT, *nia1,2* mutants and WT seedlings inoculated with *Trichoderma* and grown on the NO scavenger cPTIO. Control roots in 0.1 and 3 mM NO_3_^−^ produced reduced levels of NO. Within 2 min of *Trichoderma* application, WT plants displayed highly increased levels of NO in both 0.1 mM and 3 mM NO_3_^−^ (Fig. **5**) but the increase was slightly higher in 0.1 mM than 3 mM. That said, after 10 minutes and 24 h of *Trichoderma* incubation, the WT plants showed extremely low fluorescence (Fig. **5**). This suggests *Trichoderma* greatly induces NO in low NO_3_^−^ grown plants within a short time period. Thus, *Trichoderma* induced NO production likely plays a role in priming and induction of HATs. Assessment of NO in the *nia1,2* mutant revealed that nitrate reductase (NR) is responsible for the *Trichoderma*-induced NO production. Whilst plants grown in the presence of cPTIO displayed reduced levels of NO. We further checked the importance of *Trichoderma-elicited* NO production in the induction of HATs (*NRT 2.1 & 2.2*), total protein levels and expression of the *PR1* gene in low NO producing non-symbiotic hemoglobin over-expressing line (*nsHb^+^*), *nia1,2* and *nrt2.1* mutants. Expression of *NRT2.1* and *2.2* levels increased in WT in response to *Trichoderma*, whereas in Hb^+^ lines, *nia1,2* and *nrt2.1* mutants these two HATs were not induced (Fig. **6a,b**) suggesting that *Trichoderma* elicited NO is responsible for induction of HATs. Further, reduced protein levels were observed in *nia1,2* and Hb^+^ plants suggesting that the induction of NO_3_^−^ transporters mediated by *Trichoderma* elicited NO plays a role in increased N uptake under low NO_3_^−^ (Fig. **6c**). Furthermore, we found that *nia1,2* and Hb^+^ were unable to induce *PR1* expression under these conditions (Fig. **6d**). Taken together, these results suggest that *Trichoderma* elicited NO plays a role in overall increase in defense response under low NO_3_^−^.

**Fig. 5:**
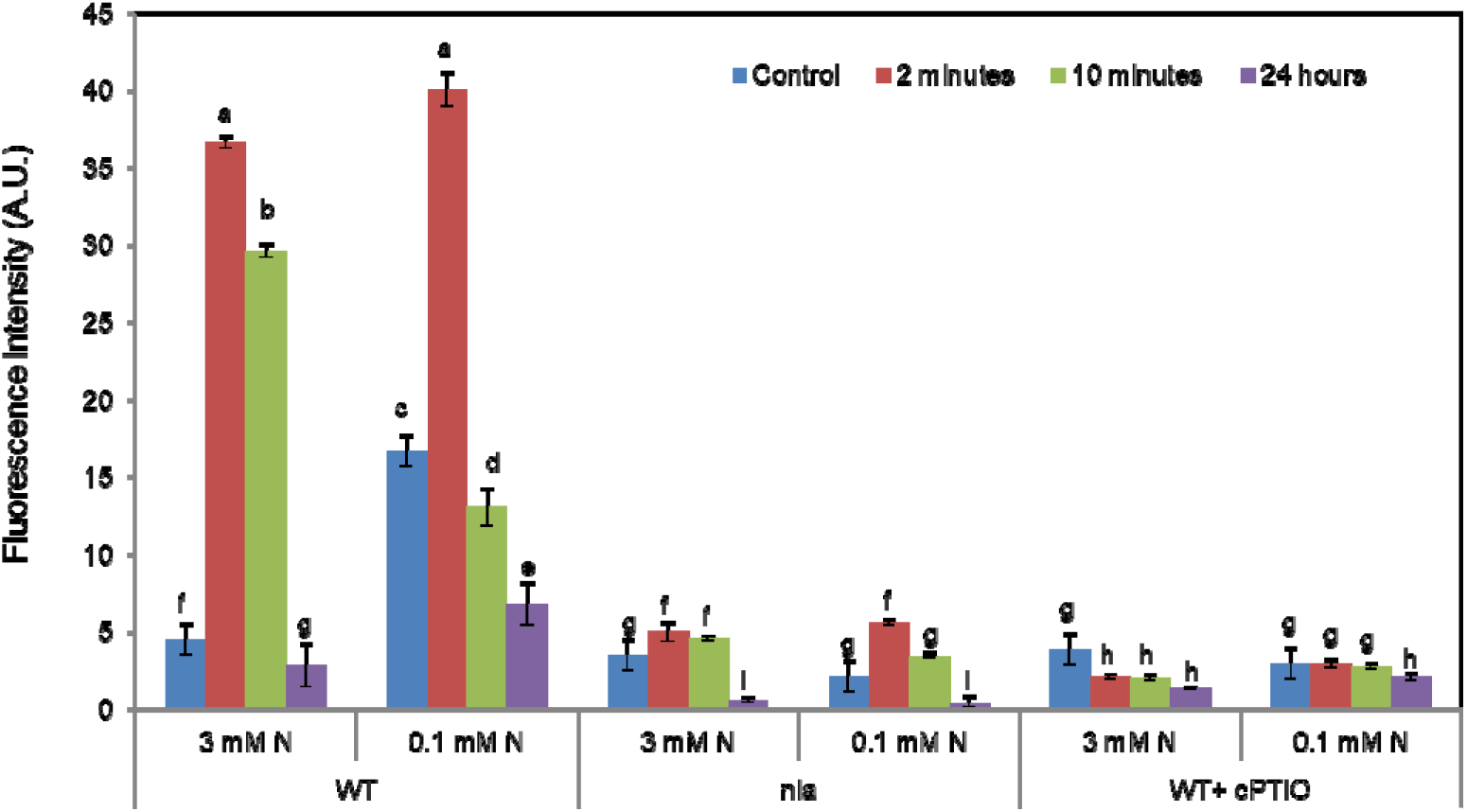
Visualization of nitric oxide by diaminofluorescein (DAF) fluorescence. Nitric oxide estimation by diaminofluorescein (DAF-FM) fluorescence under 0.1 and 3 mM NO_3_^−^ concentration in WT, *nia1,2* and cPTIO (100 μM) grown WT plants during different periods of *Trichoderma* inoculation. The experiment was performed three times independently with similar results. Data are average mean values ± SE with n=3. Statistical significance was tested by two-way ANOVA followed by Tukey’s AllPairwise Comparisons post-hoc test. The different letters above each column represent significance difference between means at p□<□0.05.

**Fig. 6:**
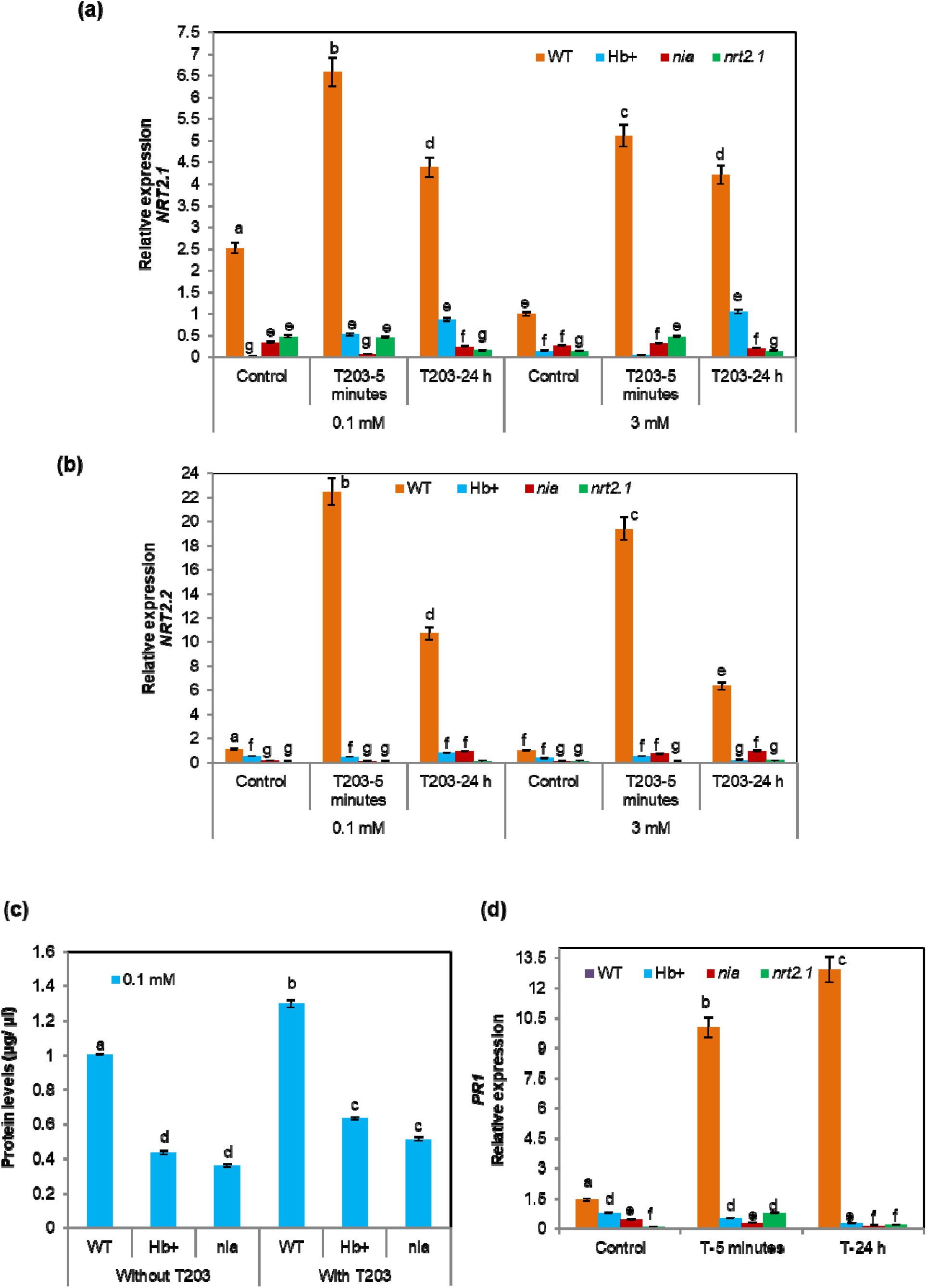
Response of the genotypes during early stages of *Trichoderma* inoculation on *NRT2.1, NRT2.2* expression, protein levels and *PR1* expression. (a) Relative *NRT2.1* expression in roots of WT, Hb^+^, *nia1,2* and *nrt2.1* plants grown under 0.1 mM and 3 mM NO_3_^−^ concentration, with and without *Trichoderma* treatment, given for 5 minutes and 24 h to the plants. Data are average mean values ± SE with n=3. Statistical significance was tested by two-way ANOVA followed by Tukey’s All-Pairwise Comparisons post-hoc test. The different letters above each column represent significance difference between means at p□<□0.05. (b) Relative *NRT2.2* expression in roots of WT, Hb^+^, *nia1,2* and *nrt2.1* plants grown under 0.1 mM and 3 mM NO_3_^−^ concentration, with and without *Trichoderma* treatment, given for 5 minutes and 24 h to the plants. Data are average mean values ± SE with n=3. Statistical significance was tested by two-way ANOVA followed by Tukey’s All-Pairwise Comparisons post-hoc test. The different letters above each column represent significance difference between means at p□<□0.05. (c) Protein levels measured in WT, Hb^+^ and *nia1,2* seedlings grown under 0.1 mM NO_3_^−^ for 15 days in vertical plates using Bradford’s assay. *Trichoderma* treatment was given for 24 h to the plants. Data are average mean values ± SE with n=3. Statistical significance was tested by one-way ANOVA followed by Dunnett’s multiple comparison test. The different letters above each column represent significance difference between means at p□<□0.05. (d) *PR1* gene expression in WT, Hb^+^, *nia1,2* and *nrt2.1* plants under 0.1 mM NO_3_^−^ plants, with and without *Trichoderma* treatment, given for 5 minutes and 24 h to the plants. Data are average mean values ± SE with n=3. Statistical significance was tested by two-way ANOVA followed by Tukey’s All-Pairwise Comparisons post-hoc test. The different letters above each column represent significance difference between means at p□<□0.05.

We next investigated the role of NO in LAR and SAR development in the presence of *Trichoderma* using NO mutants. Under low NO_3_^−^, WT plants displayed a stronger LAR response than either the Hb^+^ lines or the *nia1,2* mutant. Furthermore, reduced bacterial growth and increased *PR1* gene expression was observed in WT compared to either the Hb^+^ lines or the *nia1,2* mutant (Fig. **7 a,b,c**). Similarly, the SAR response was also more compromised in the WT than in the Hb+ lines and the *nia1,2* mutant ((Fig. **7 d,e,f**). As for LAR, a similarly reduced bacterial number and increased *PR1* gene expression was observed in WT in comparison to the Hb^+^ lines and the *nia1,2* mutant

**Fig. 7:**
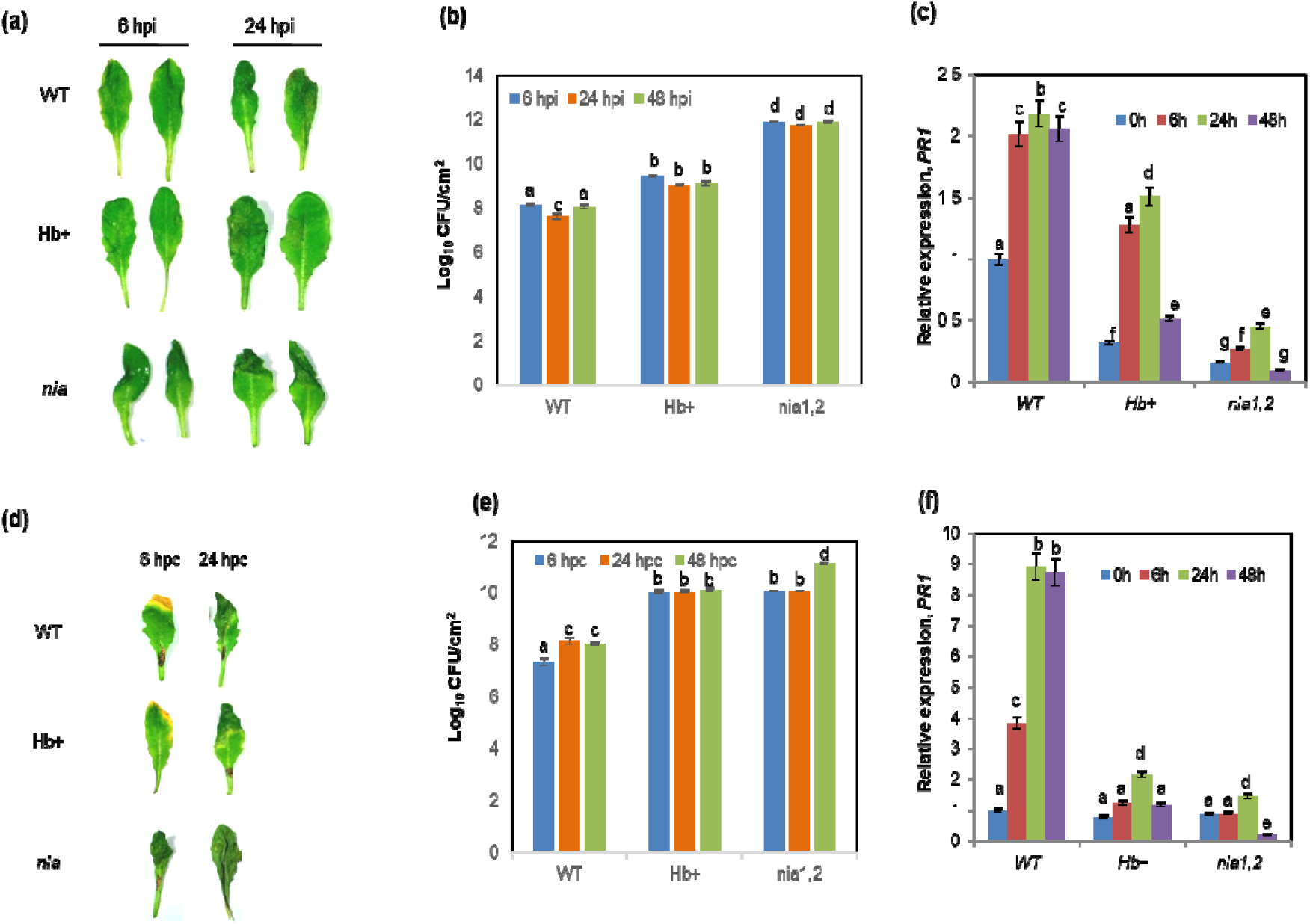
Response of WT and NO mutants (Hb^+^ lines and *nia1,2*) grown under 0.1 mM NO_3_^−^ during LAR and SAR. In LAR response (a) HR phenotype in inoculated leaves of WT, Hb^+^ and *nia1,2* plants grown under 0.1 mM NO_3_^−^ concentration at 0 and 24 hpi. (b) In planta bacterial growth (log CFU) at 6, 24 and 48 hpi. Data are average mean values ± SE with n=3. Statistical significance was tested by one-way ANOVA followed by Dunnett’s multiple comparison test. The different letters above each column represent significance difference between means at p□<□0.05. (c) Relative *PR1* expression in infiltrated leaves during localized infiltration. Data are average mean values ± SE with n=3. Statistical significance was tested by two-way ANOVA followed by Tukey’s All-Pairwise Comparisons post-hoc test. The different letters above each column represent significance difference between means at p□<□0.05. In SAR response (d) HR phenotype in inoculated and uninoculated leaves of WT, Hb^+^ and *nia1,2* plants grown under 0.1 mM NO_3_^−^ concentration at 0 and 24 hours post secondary challenge (e) Bacterial population represented by log CFU at 6, 24 and 48 hpc. Data are average mean values ± SE with n=3. Statistical significance was tested by one-way ANOVA followed by Dunnett’s multiple comparison test. The different letters above each column represent significance difference between means at p□<□0.05. (f) Relative *PR1* gene expression in infiltrated leaves post secondary challenge. Data are average mean values ± SE with n=3. Statistical significance was tested by two-way ANOVA followed by Tukey’s All-Pairwise Comparisons post-hoc test. The different letters above each column represent significance difference between means at p□<□0.05.

### ROS is a component of the *Trichoderma*-induced increased resistance via SAR

Both NO and ROS are involved in the plant resistance response hence we next investigated the role of ROS. The distal leaves of 0.1 mM NO_3_^−^ grown WT plants in the presence of *Trichoderma* showed increased H_2_O_2_ production in comparison to 0.1 mM NO_3_^−^ grown distal leaves (Fig. **8a**). Also, the level of H_2_O_2_ was more in 3 mM NO_3_^−^ grown distal leaves of WT plants in the presence of *Trichoderma* in comparison to untreated plants (Fig. **8a**). This is probably due to the suppression of catalase activity by SA. This suggests that *Trichoderma* inoculation can enhance H_2_O_2_ levels in distal leaves during SAR, whereas, both the *Pst* inoculated leaves from WT grown under 3 mM NO_3_^−^ and WT plants in the presence of *Trichoderma* displayed higher H_2_O_2_ production (Fig. **8a**). This clearly suggests that *Trichoderma* plays a role in inducing H_2_O_2_ in low NO_3_^−^ fed WT plants during SAR.

**Fig. 8:**
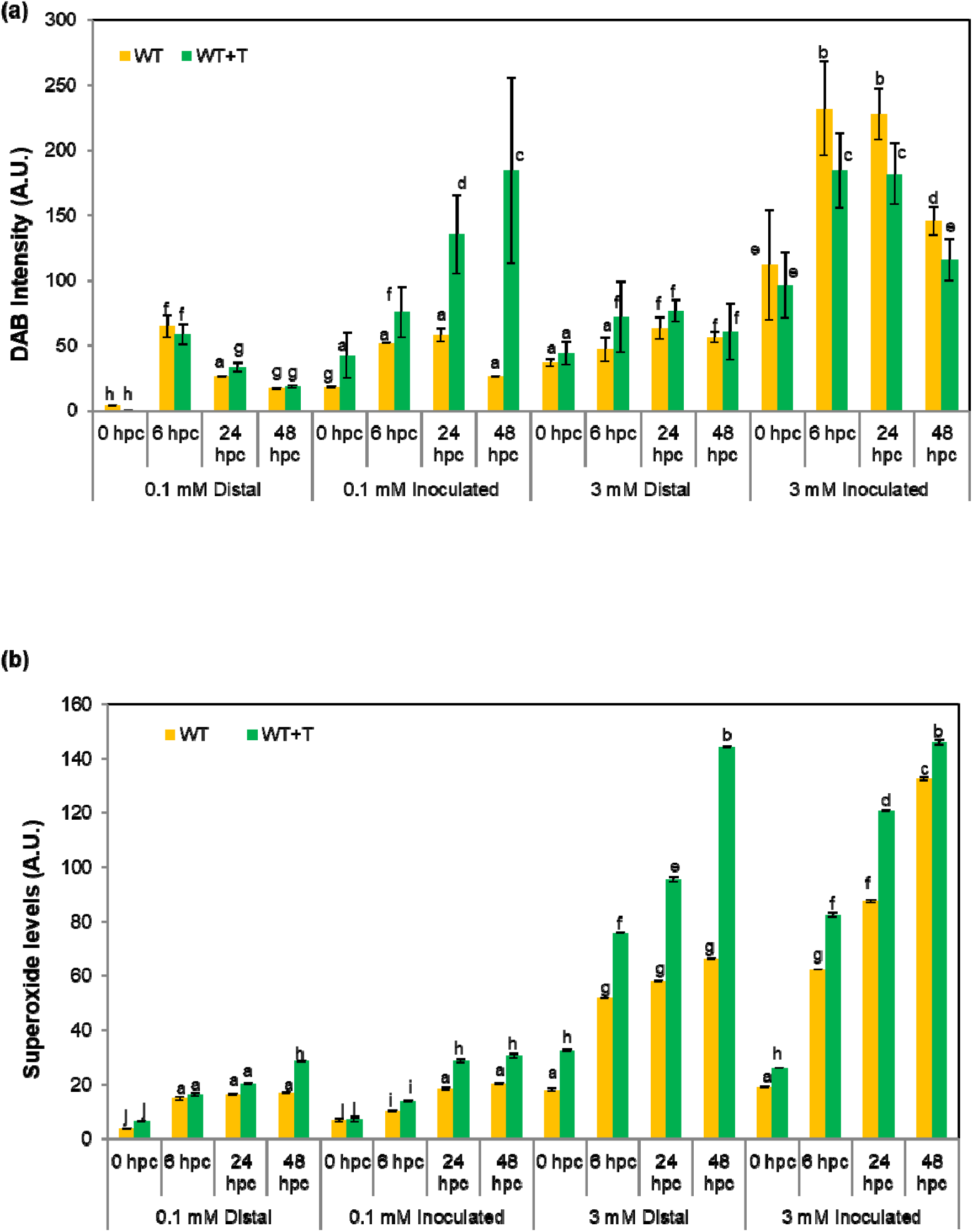
Detection of ROS by measuring H_2_O_2_ and O_2_^−.^ in 0.1 mM and 3 mM NO_3_^−^ WT and *Trichoderma* grown WT plants during SAR response. (a) Quantification of DAB staining used to measure H_2_O_2_ levels in the inoculated leaves and distal leaves post challenge inoculation at 0, 6, 24 and 48 hpc is shown as intensity per stained area from ten leaves per treatment measured using ImageJ in arbitrary units with the mean values ± SE. Column graph represents the DAB intensity calculated as stained leaf area compared with the total surface of leaf analyzed in terms of % stained area upon the % control area by ImageJ (ver 3.2). The control leaves were completely decolorized using bleaching agent and then the area was measured using ImageJ and % was calculated with total area and the result obtained was termed % control area. Statistical significance was tested by two-way ANOVA followed by Tukey’s All-Pairwise Comparisons post-hoc test. The different letters above each column represent significance difference between means at p□<□0.05. (b) Quantification of NBT staining used to measure O_2_^−^ levels in the inoculated leaves and distal leaves post challenge inoculation at 0, 6, 24 and 48 hpc shown as intensity per stained area from ten leaves per treatment measured using ImageJ in arbitrary units with the mean values ± SE. Similar method was used to measure % stained area for NBT stained images w.r.t control leaves. Statistical significance was tested by two-way ANOVA followed by Tukey’s All-Pairwise Comparisons post-hoc test. The different letters above each column represent significance difference between means at p□<□0.05.

O_2_^−.^ is a key player in cell death (Fig. **8b**). It was observed that, there was increased O_2_^−.^ in inoculated leaves of both 0.1 and 3 mM NO_3_^−^ fed WT and *Trichoderma* treated WT plants, but the increase was much higher in the presence than in the absence of *Trichoderma*. This suggests that there was a rapid oxidative burst (HR) which offers resistance to the plants in response to *Trichoderma* treatment. Taken together, the coordinated interplay of H_2_O_2_, O_2_^−.^ and NO leads to the HR associated cell death that greatly improves the LAR and SAR responses of low N-stress plants.

### *Trichoderma* induces defense genes during SAR in low N-fed plants

We next checked the expression levels of defense related genes (*PAL1, PR1, PR2* and *PR5*) during SAR. In both 0.1 mM and 3 mM NO_3_^−^ grown, *Pst* inoculated WT plants, *PAL1* transcripts were highly induced in all time points, but, the distal leaves showed *PAL1* induction only until 6 hpc, and this dramatically declined at later time points (Fig. **9a**). In the presence of *Trichoderma* WT plants grown under 0.1 mM and 3 mM NO_3_^−^ regimes, showed even greater enhancements in the levels of *PAL1* transcripts) in inoculated as well as distal leaves until 48 hpc. A similar trend was observed in the *PR1* (Fig.**9b**), *PR2* (Fig. **9c**) and *PR5* (Fig. **9d**) expression profiles, which displayed elevation following *Trichoderma* application. Of the four defense genes examined, the *PR1* gene displayed the highest expression levels in plants grown in the presence of *Trichoderma* suggesting that *Trichoderma* might induce SA levels to a greater extent during SAR.

**Fig. 9:**
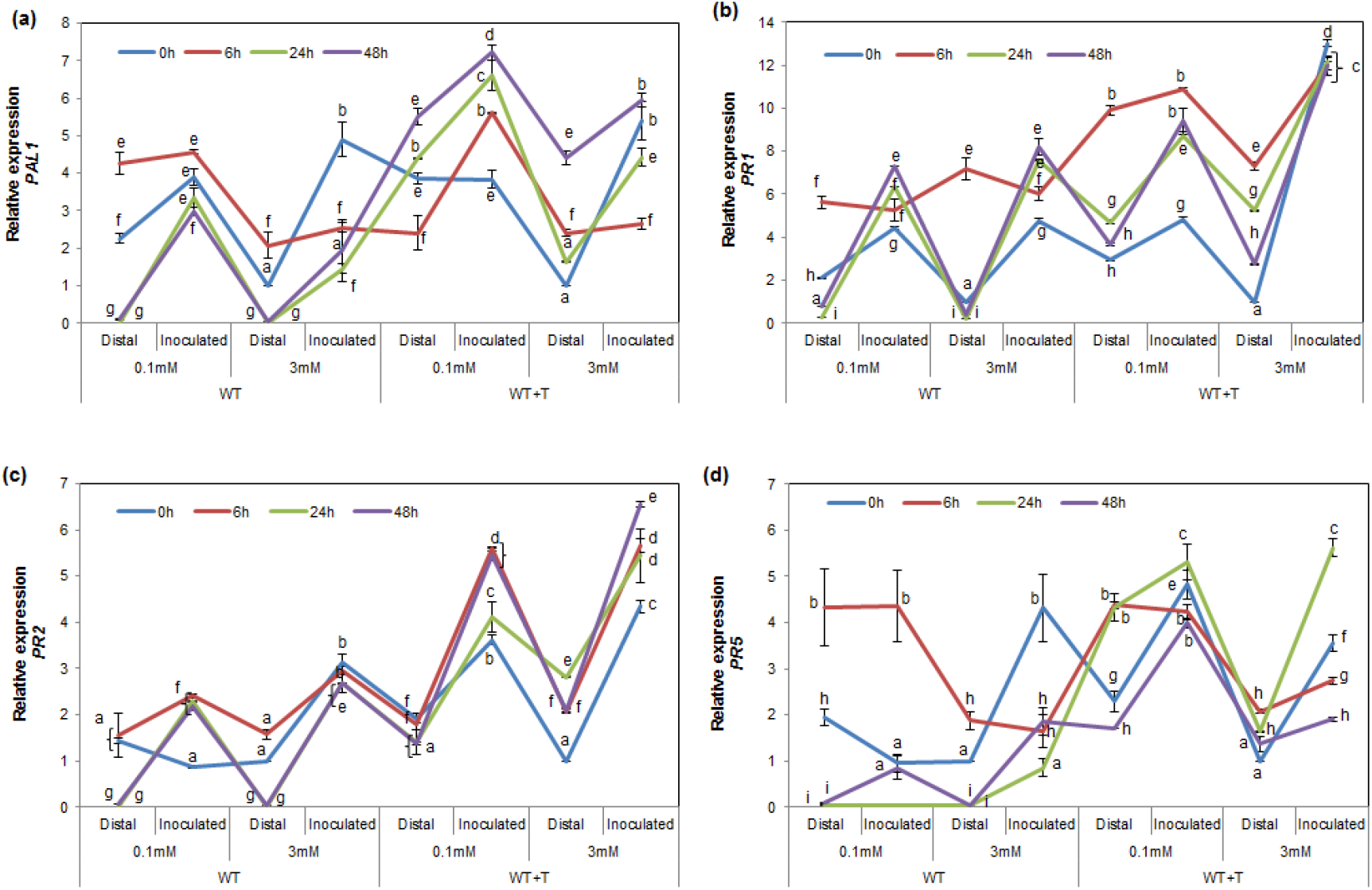
Expression profiles of defense related genes during SAR. Relative expression of defense related genes in WT and WT+T grown under low (0.1 mM) and optimum (3 mM) NO_3_^−^ concentration post-secondary challenge of *Pst*DC3000 in both inoculated and distal leaves (a) Relative expression of *PAL1* (b) *PR1* (c) *PR2* (d) *PR5*. For all the target genes, fold expression values are means (n=3) ± SE. Statistical significance was tested by two-way ANOVA followed by Tukey’s All-Pairwise Comparisons post-hoc test. The different letters above each column represent significance difference between means at p□<□0.05.

### *Trichoderma* enhances expression of SAR-mediated regulatory genes

Having established the link between *Trichoderma* and SAR we next checked whether *Trichoderma* can induce the SAR response via induction of regulatory genes such as *DIR1, NPR1, SARD1 and TGA3*. The lipid transfer protein, DEFECTIVE IN INDUCED RESISTANCE1 (*DIR1*) is a key mobile component of SAR response (Maldonaldo *et al*. 2002) involved in long-distance translocation from local to distant leaves (Carella *et al*. 2015, Champigny *et al*. 2013). It was found that, *DIR1* induction took place only in the initial time points in inoculated and distal leaves of both 0.1 and 3 mM NO_3_^−^ grown plants (Fig. **10a**). However, in the presence of *Trichoderma*, a slightly increased *DIR1* expression was observed at all time points (Fig. **10a**). Upon perception of SAR mobile signals, *Non-Expresser of Pathogenesis-Related gene1* (*NPR1*) activates defense in challenged plants (Cao *et al*. 1997). We found a similar trend in the *NPR1* expression profile as we did for *DIR1* (Fig. **10b**). Moreover, upon *Trichoderma* treatment, the levels of *NPR1* inoculated and distal challenged leaves gradually increased in both 0.1 and 3 mM NO_3_^−^ grown (Fig. **10b**). Following this we checked the expression of *SAR DEFICIENT 1* (*SARD1*) a pathogen-induced transcription factor (Zhang *et al*. 2010) and a key regulator of *Isochorismate Synthase 1* (*ICS1*) and SA synthesis (Wang *et al*. 2011). A remarkably stronger induction of *SARD1* expression levels in *Trichoderma* treated distal leaves of 0.1 mM NO_3_^−^ grown plants at 6 hpc (~82 fold) revealed that is a potential inducer of SA biosynthesis (Fig. **10c**). Another important regulatory gene, *TGA3*, is an *NPR1*-interacting protein (NIP) as well as a critical component in the SA signaling mechanism. This gene was induced in all treatments in response to *Trichoderma* (Fig. **10d**) highlighting the important *Trichoderma*-mediated promotion of SA signaling pathways.

**Fig. 10:**
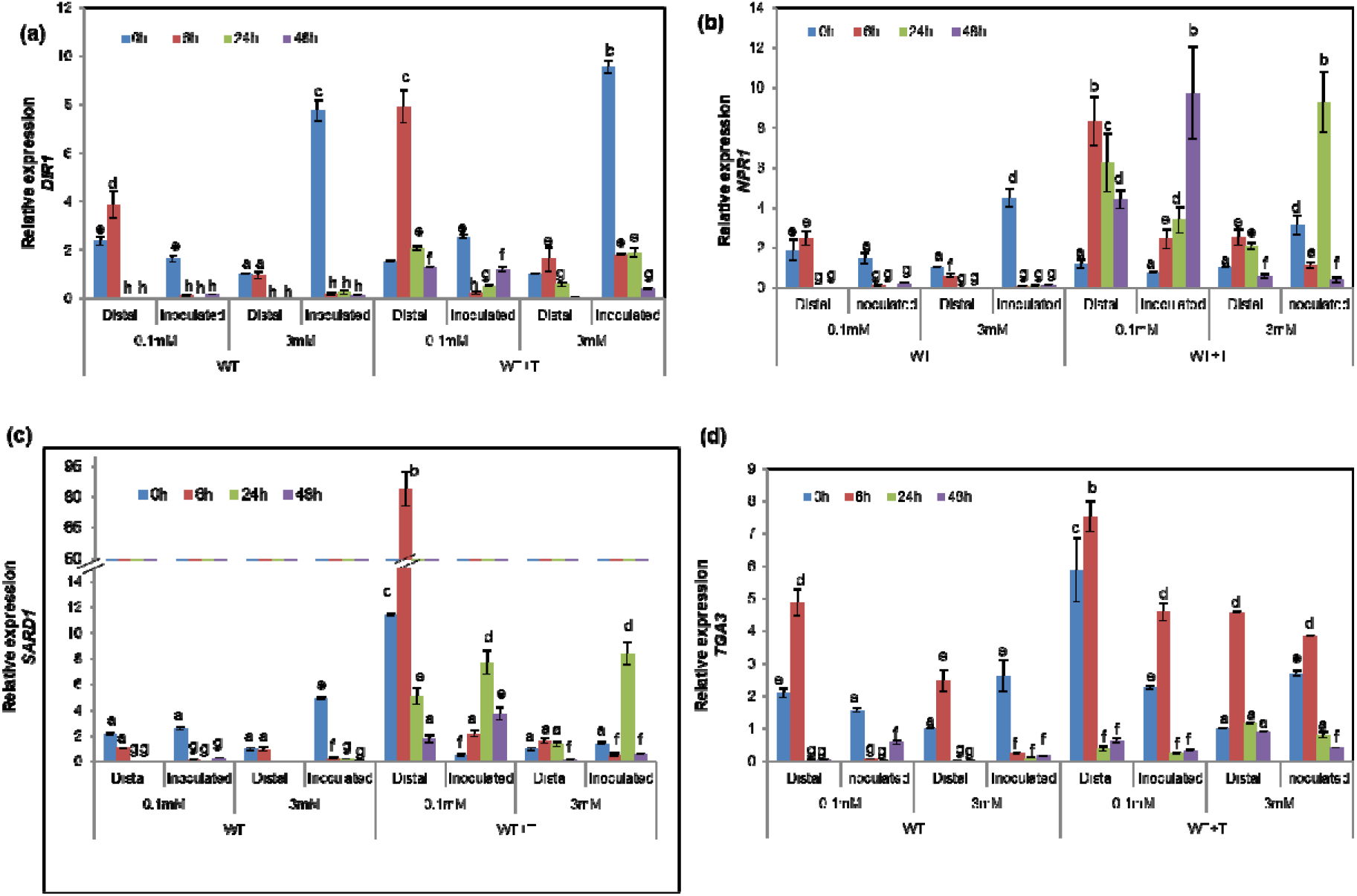
Expression profiles of regulatory SAR genes. Relative expression of SAR regulatory gene in WT and WT+T grown under low (0.1 mM) and optimum (3 mM) NO_3_^−^ concentration post-secondary challenge of *Pst*DC3000 in both inoculated and distal leaves. Relative expression of (a) *DIR1* (b) *NPR1* (c) *SARD1* (d) *TGA3*. For all the target genes, fold expression values are means (n=3) ± SE. Statistical significance was tested by two-way ANOVA followed by Tukey’s All-Pairwise Comparisons post-hoc test. The different letters above each column represent significance difference between means at p□<□0.05.

### nrt2.1 and npr1 mutants are compromised in LAR and SAR responses

Having clearly demonstrate the link between N nutrition and defense we next decided to characterize LAR in *nrt2.1* and *npr1* mutants in response to virulent *PstDC3000*. The 0.1 mM NO_3_^--^ fed *nrt2.1* and *npr1* mutants developed more severe symptoms than 0.1 mM NO_3_^−^-fed WT plants (Fig. **11a** 0.1 mM panel). An increased CFU count was observed in 0.1 mM NO_3_^−^ fed WT and *nrt2.1* and *npr1* mutants at 24 and 48 hpi in comparison to bacterial numbers in 3 mM NO_3_^−^ grown plants in which overall bacterial count of WT was less but a significant increase in CFU was observed in *nrt2.1* and *npr1* mutants (Fig.**11b**). This suggests that optimum NO_3_^−^ concentration plays an important role in resistance response towards virulent *Pst* DC3000.

**Fig. 11:**
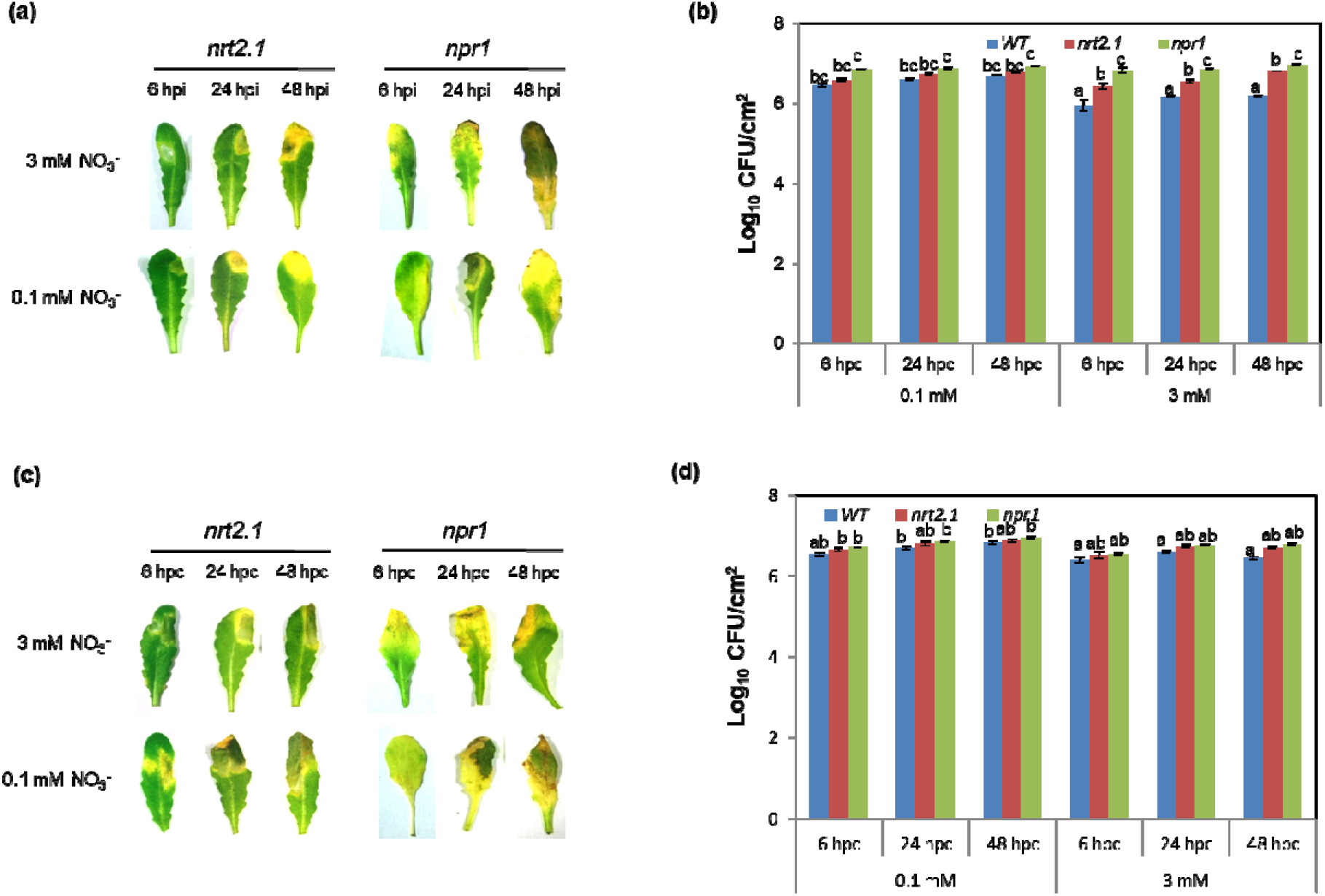
LAR and SAR response in *nrt2.1* and *npr1* mutants to virulent *Pst* DC3000. (a) LAR disease symptoms to virulent Pst DC3000 in *nrt2.1* and *npr1* mutant in 0.1 and 3 mM nitrate (b) Bacterial growth in WT, *nrt2.1* and *npr1* mutant (c) Phenotype during SAR response in *nrt2.1* and *npr1* mutant in 0.1 and 3 mM nitrate (d) Bacterial growth in WT, *nrt2.1* and *npr1* mutant during SAR response.

Similarly, we studied the SAR response in the WT, *nrt2.1* and *npr1* plants. 0.1 mM NO_3_^−^ grown *Pst*-challenged WT plants show more prominent necrotic and discolored lesions spread to half of the leaf (Fig. **2a**) confirming susceptible symptoms (yellow specks). On the other hand, the *nrt2.1* and *npr1* mutants showed even more severe disease symptoms – namely extensive chlorosis and necrosis (Fig. **11c**) under low NO_3_^−^ growth conditions. Our results suggest that, 3 mM NO_3_^−^ grown plants proved more resilient to secondary challenge than 0.1 mM NO_3_^−^ grown plants. Bacterial populations were also increased in the *nrt2.1* and *npr1* mutants (Fig. **11d**) suggesting that *NRT2.1* and *NPR1* plays an important role in increasing defense mediated by *Trichoderma*.

### Salicylic acid pathway is a part of enhanced plant resistance mediated by *Trichoderma* under low N

Examination of SA levels revealed that *Trichoderma* presence accelerated total SA levels in 0.1 mM distal leaves, in comparison to 3 mM distal leaves. There was significant increase in SA levels observed in inoculated and distal leaves of *Trichoderma* grown plants, in comparison to SA levels in WT plants (Fig 12a). We sought to further confirm role of SA in *Trichoderma* increased SAR, hence *nahG* plants were challenged in the presence or absence of *Trichoderma*. An intense chlorotic lesion was evident in both 0.1 and 3 mM grown *nahG* plants in response to challenge inoculation while *Trichoderma* grown *nahG* plants when challenged they defended much better evidenced by decreased chlorotic lesions and reduced bacterial numbers (Fig. **12 b,c**). Surprisingly, *Trichoderma* grown *nahG* plants showed slightly enhanced *PR1* transcript levels (Fig. **12d**).

**Fig. 12:**
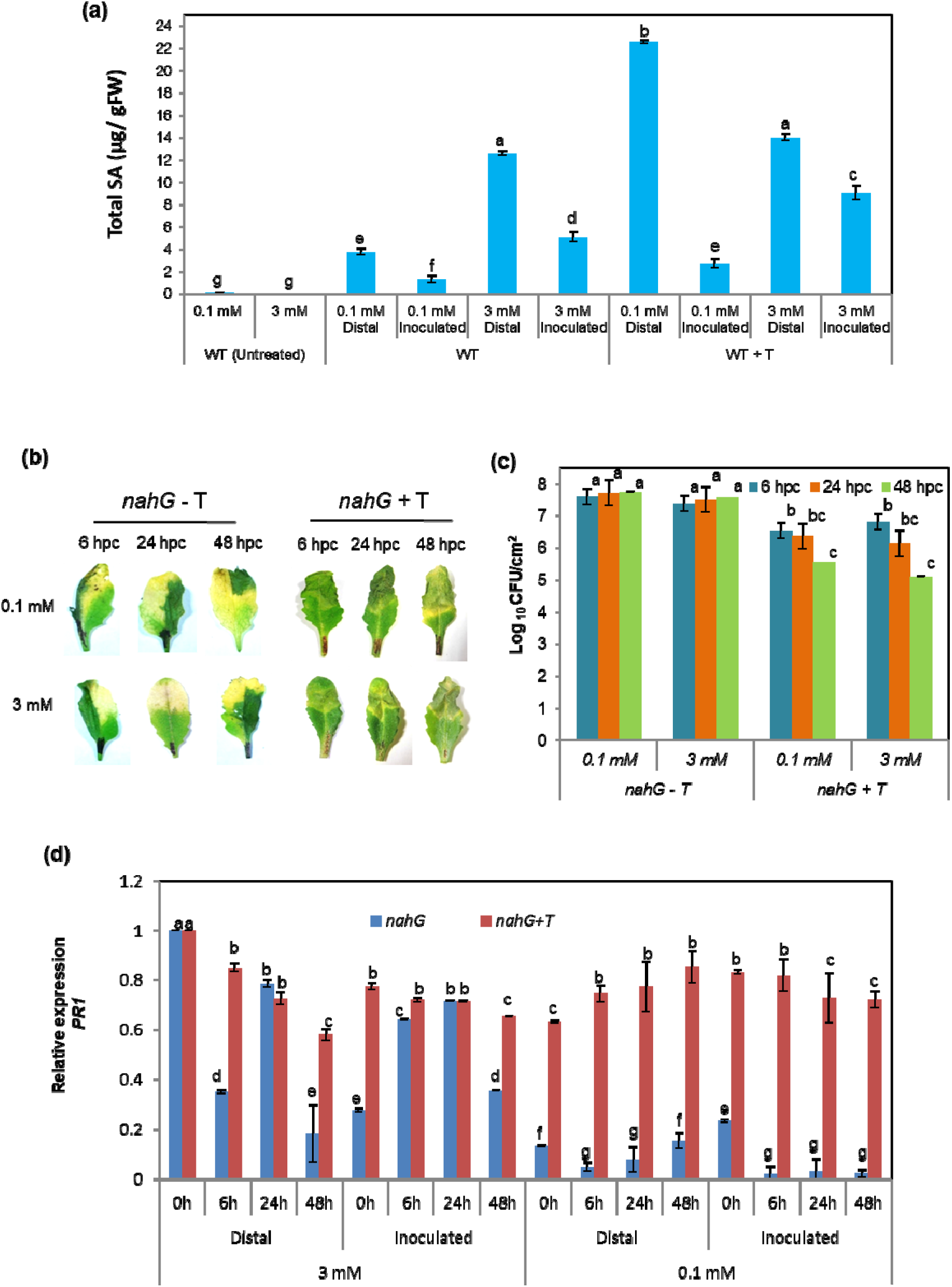
SA Accumulation during SAR, and response of *nahg* in the presence and absence of *Trichoderma* during SAR. (a) Total SA levels (μg g^−1^ fresh weight) were determined in untreated WT leaves (without *Trichoderma* and *Pst* treatment) and in inoculated and distal leaves of 0.1 mM and 3 mM nitrate-fed WT plants post challenge inoculation (24 hpc) with and without *Trichoderma*. Data are average mean values ± SE with n=3. Statistical significance was tested by two-way ANOVA followed by Tukey’s All-Pairwise Comparisons post-hoc test. The different letters above each column represent significance difference between means at p□<□0.05. (b) HR phenotype in inoculated and un-inoculated distal leaves of *nahg* mutants grown under 0.1 and 3 mM nitrate concentration post-secondary challenge at 6, 24 and 48 hpc (c) Bacterial number represented by log CFU at 6, 24 and 48 hpc from the inoculated leaves post secondary challenge (d) *PR1* gene expression in inoculated and uninoculated leaves of *nahG* mutants grown under 0.1 and 3 mM nitrate, with and without *Trichoderma* treatment post-secondary challenge. Data are average mean values ± SE with n=3. Statistical significance was tested by two-way ANOVA followed by Tukey’s All-Pairwise Comparisons post-hoc test. The different letters above each column represent significance difference between means at p□<□0.05.

## Discussion

Nitrogen availability and supply can severely impact growth and development of plants (Walker *et al*. 2001; Landrein *et al*. 2018). N deficiency can cause chlorosis, which can impact photosynthesis and overall energy demand for growth and defense with investment in plant resistance-related compounds being drastically constrained under limiting nitrogen supply (Dietrich *et al*. 2004). This suggest a possibility that increased N uptake under limited supply might boost defense. Previously it was shown that N levels have been shown to affect the synthesis of constitutive defences based on secondary metabolites such as alkaloids (Stout *et al*., 1998) and (poly) phenolics under varying N regimes (Johnson *et al*., 1987). Since N is also important for synthesis of various secondary metabolites, severe depletion of N can also impact defense related pathways. Hence, plants may not be able to activate adequate defense pathways for tolerance or resistance (Snoeijers *et al*. 2000). Ward *et al*. (2010) have previously shown that plant reconfigure their metabolism in response to N supply. Plants take up N in the form of NH_4_^+^ or NO_3_^−^ or a combination of both. Different N forms differentially effect various free radicals such as NO and ROS (Wany *et al*. 2018; Gupta *et al*. 2013). Ammonium uptake and assimilation is less costly to the plants in comparison to NO_3_^−^ but excess of NH4^+^ can cause toxic effects to the plants (Boudsocq *et al*. 2012; Liu *et al*. 2017), hence, many plants preferentially use NO_3_^−^ as N source. NO_3_^−^ nutrition can also enhance plant defence via increased generation of NO, polyamines and SA (Gupta *et al*. 2013; Fagard *et al*. 2014). Under NH_4_^+^ nutrition plants compromise in resistance due to reduced NO, increased sugars, aminoacids and enhancement of γ-aminobutyric acid (GABA) levels which may be source of nutrition for pathogens (Gupta *et al*. 2013) Hence, here we chose to check the plant defense response under NO_3_^−^ nutrition rather than under NH_4_^+^.

Plants grown on 0.1 mM NO_3_^−^ showed reduced growth (Fig. **S3**) and morphological parameters (Fig. **S3**), suggesting that the supplied 0.1 mM NO_3_^−^ is insufficient for optimal growth of plants and consequently would likely reduce the available resources which could be allocated to defense. In our experiments, 0.1 mM NO_3_^−^ grown plants were compromised in both LAR and SAR (Fig. **1****,2**) responses, suggesting that NO_3_^−^ is required for better defense. Plants grown on low NO_3_^−^ produced less SA (Fig. **12a**), further supporting the notion that NO_3_^−^ is needed for SA biosynthesis. Since plants need N for growth and disease resistance, increasing their N use efficiency may be an effective manner to improve their resistance. Certain types of *Trichoderma* aid in nutrient absorption leading to increased growth and enhanced plant defense (Brotman *et al*. 2010). It was previously demonstrated that supplementation of plants with *Trichoderma asperelloides* enhances plant growth (Brotman *et al*. 2012), and protects against abiotic and biotic stressors-moreover, It was, moreover, demonstrated to induce systemic resistance responses (Contreras-Cornejo *et al*. 2016; Brotman *et al*. 2012). *Trichoderma* induced increased growth has been varyingly attributed to auxin and ethylene (Garnica-Vergara *et al*. 2016) and the induction of genes involved in carbon and N metabolism (Domínguez *et al*. 2016). However, there are hardly any reports concerning the mechanism(s) underlying *Trichoderma*-mediated plant growth and defense improvement under N starvation. In the current study, we unraveled the mechanism of *Trichoderma*-induced plant growth and defense under low N. Hence, in this current work, we studied the impact of *Trichoderma* on enhancing N uptake and supporting both LAR and SAR responses under low NO_3_^−^ stress.

The enhanced growth of plants grown in the presence of *Trichoderma* is due to an increased NO_3_^−^ uptake (Fig. 3a) which was also evidenced by increased expression of *NRT2.1, NRT2.2* (Fig. **4a,b,c,d,e**) and increased total cellular protein levels (Fig. **4f**). *NRT2.1* is the main HAT, localized at the plasma membrane. Previously, it was shown that these transporters become active during N starvation and are severely inhibited when reduced nitrate sources such as glutamine or ammonium are provided (Dechorgnat *et al*. 2012). Similarly, *NRT2* is also induced under low N (Dechorgnat *et al*. 2012). In response to *Trichoderma* treatment, a rapid induction of *NRT2.1, NRT2.2* transcripts in both plants grown with 0.1 and 3 mM NO_3_^−^ was observed during *Pst* inoculation. *NRT2.1* involvement was further evidenced by the fact that the *nrt2.1* mutant produced less protein under low N and even the addition of *Trichoderma* was unable to increase protein content in this mutant (Fig. **4f**). Furthermore, this mutant become highly susceptible to LAR and SAR under low N (Fig. **11**). It thus appears reasonable to assume that *Trichoderma* can increase N uptake and by this means enhance resistance

Hence, in further experiments, we focused on the LAR and SAR responses under low and optimum NO_3_^−^ in the presence or absence of *Trichoderma*. During local *Pst* infection, the plants display a LAR response, and the systemic/distal leaves induce a SAR response. Both are critically important for plant defense against pathogens.

In 3 mM NO_3_^−^-fed WT plants, the defense response after the secondary challenge was also more rapid, robust and even longer-lasting till 5 days post challenge (data not shown) whereas, 0.1 mM NO_3_^−^ fed WT plants showed disease symptoms suggesting that NO_3_^−^ concentration plays a key role in the development of both LAR and SAR (Fig. 1a, **2a** and **S1b**).

One of the features of *Trichoderma* is the induction of short-term spikes of NO This molecule play an important role in induction of plant defense responses (Gupta *et al*. 2014). We suspected the role of NO in activating genes of these transporters. Indeed, *Trichoderma* induced expression of *NRTs* are most likely mediated by short term increase in NO upon *Trichoderma* inoculation. NR-dependent NO elicited by *Trichoderma* is probably responsible for the increased expression of HATs since our experiments revealed that the Hb^+^ lines and the *nia1,2* mutant, were unable to induce HATs and showed decline in protein levels even in the presence of *Trichoderma* (Fig. **6c**). *Trichoderma* trigger the SA-dependent SAR pathway (Pieterse *et al*. 2014), they induced ROS involved in the plant’s resistance response against many biotic stressors (Asmawati *et al*. 2017). They additionally plays an important role in hypersensitive cell death together with NO (Durner and Klessig 1999, Dorey *et al*. 1999). Among ROS, H_2_O_2_ is the most stable form which plays an important role as a signal transducer in the plant cell death process (Pieterse *et al*. 2014), and acts as a key modulator of NO in triggering cell death. As shown in Fig. **8a** *Trichoderma* treated low N fed plants displayed increased H_2_O_2_ levels, thus, enabling the initiation and the establishment of SAR. Superoxides (O_2_^−.^) are mainly produced via mitochondrial electron transport and NADPH oxidases during stress plays a role in plant defense (Torres *et al*. 2002, 2005). Earlier, it was shown that the extracellular elicitors isolated from *Trichoderma viridae* also induces O_2_^−.^ levels (Calderon *et al*. 1994). Here, we also found that in the presence of *Trichoderma*, 0.1 mM NO_3_^−^ grown *Pst* inoculated plants show higher O_2_^−.^ levels than in the absence of *Trichoderma* (Fig.**8b**), suggesting that it can enhance O_2_^−.^ production during infection which can aid in defense. Consequently, the expression of defense marker genes such as *PR1*, *PR2, PR5* and *PAL1* (Fig. **9**) were also highly induced in the presence of *Trichoderma* in low NO_3_^−^-fed plants suggesting that *Trichoderma*-mediated ROS andNO along with increased N are probably responsible for higher induction of these genes. Martinez-Medina *et al*. (2013), reported several *Trichoderma* strains are known to induce systemic responses by acting as a “short circuit” in plant defense signaling. *PAL1* is an important marker gene in SA mediated defense (Kim and Hwang 2014) and accumulates in cells undergoing HR (Dorey *et al*. 1997) and is thought to be essential for local and systemic resistance (Delaney *et al*. 1994). The fact that *PAL1* levels were increased by *Trichoderma* is intriguing. The SAR response is associated with a specific set of SAR genes encoding pathogenesis related (PR) proteins (Pieterse *et al*. 1996). In our study these *PR* genes are activated and consequently accumulate during SAR (Fig.**9b,c,d**) in keeping with the earlier findings of Brotman *et al*. (2012) and Pieterse *et al*. (1996).

In our study, the SAR regulatory genes *DIR1, NPR1* and *TGA3* are induced in the presence of *Trichoderma* under low N stress. The *npr1* mutant is compromised in LAR and SAR in the presence of *Trichoderma* suggesting a role for this gene in enhancing defense. In the *nahG* transgenic line, defense responses are slightly enhanced in the presence of *Trichoderma* in spite of reduced SA levels, suggesting that apart from SA other factors such as increased NO and ROS are probably responsible for defense in this mutant. Overall, our study demonstrated that optimum N is required for both LAR and SAR. *Trichoderma* can enhance N uptake via modulating N transporters and via eliciting short term NO under low NO_3_^−^ nutrition.

The enhanced N uptake plays a role in enhancing SA levels and defense gene expression in local and distal leaves to increase overall plant defense (Fig. **13**). These defense responses are neither activated in *npr1,, nrt2.1*, or *nia1,2* mutants; nor in the *nahG* and *Hb^+^* transgenic lines providing corroborative evidence that *Trichoderma* mediated enhanced resistance under low NO_3_^−^ involves synergistic roles of NO, ROS and SA. Further work is needed on how this mechanism works in key crop plants. Understanding the role of *Trichoderma* in improving nitrogen use efficiency (NUE) in response to different pathogens which have diverse strategies can help in improving plant resistance to pathogens and improving plant productivity when limited N available. N is important for plant defense as well as growth hence using beneficial *Trichoderma* can be a great tool to confer multifaceted benefits to the plants and improve crop resilience under reduced N availability

**Fig. 13:**
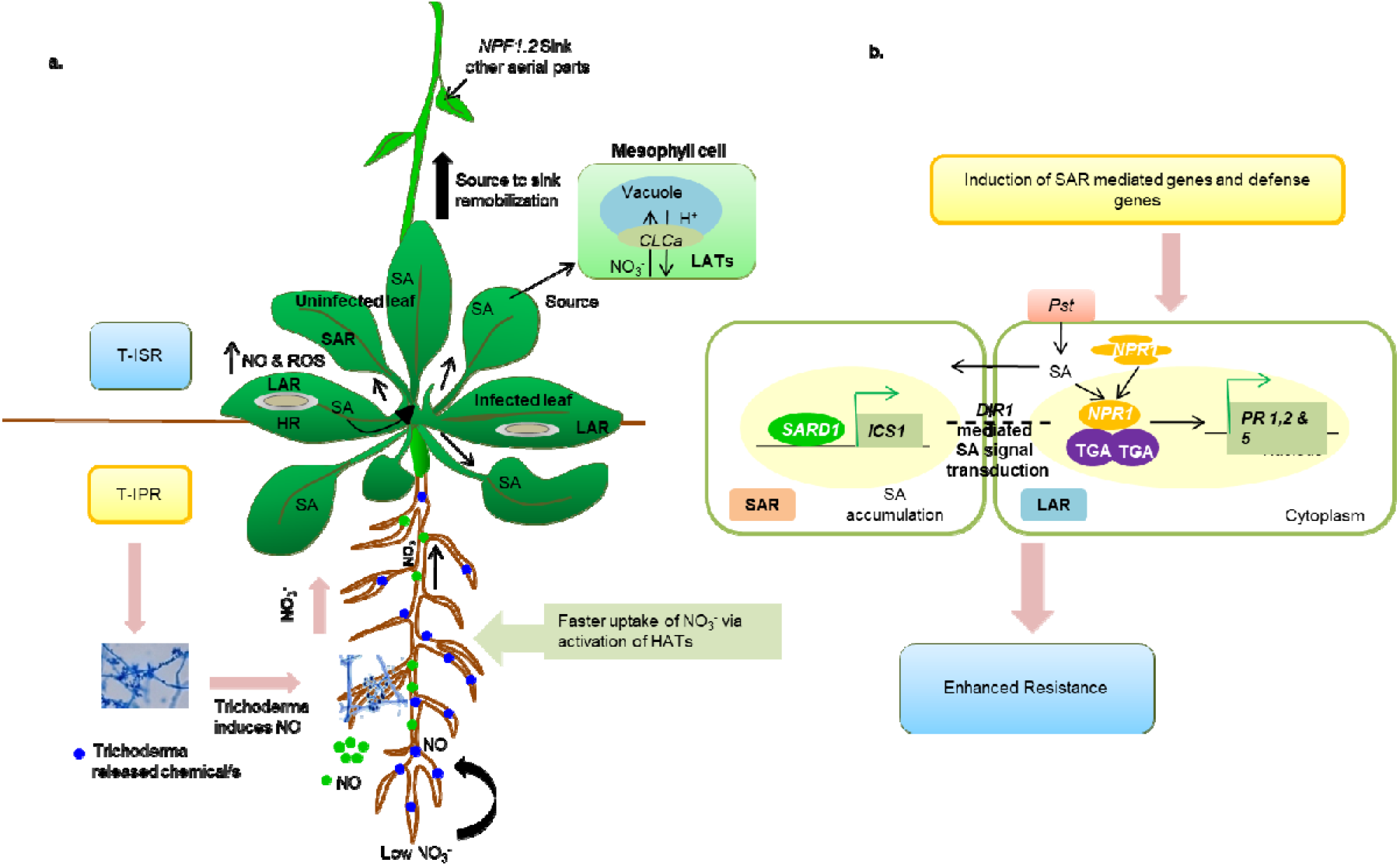
A model depicting mechanism of *Trichoderma* induced systemic response (T-ISR) under low nitrate conditions during pathogen infection in Arabidopsis. (a) After root colonization, *Trichoderma* secretes growth promoting elicitors (blue dots), and induces priming response (T-IPR) by eliciting short term nitric oxide (NO; green dots) which facilitate faster nitrate uptake by activating HATs (*NRT2.1, 2.2* and *2.4*) in the roots. The HATs in turn activates the vacuolar LATs (*CLCa* and *NPF1.2*) in the mesophyll cells allows source to sink re-mobilization of available nitrate from roots to other aerial parts. During primary inoculation (local pathogen attack; *avr*Rpm1), NO and ROS signals both are produced during hypersensitive response (HR), along with salicylic acid (SA), but their basal levels are greatly enhanced due to *Trichoderma* treatment. Moreover, the *Trichoderma* induced short term NO produced during priming and HR, plays a role in faster nitrate uptake by the HATs in low N stress plants. *Trichoderma* induced SA (produced during local infection) gets rapidly translocated to the other uninfected distal parts of the plant and pre-programs the stressed plant for subsequent pathogen attack. (b) *Trichoderma* activates PR proteins and cause SA accumulation in locally infected leaves during T-ISR. These signals are transported to the other part of the plant by activating a set of regulatory genes (*DIR1, NPR1, SARD1, TGA3*) involved in SAR response. The SA signal transduction mediated by *DIR1 NPR1 and TGA3* was evidenced by their induced expression in systemic leaves in the presence of *Trichoderma*. Moreover, induced *SARD1* expression activates the SA biosynthetic genes (*ICS1*) in the systemic leaves. All these genes are involved in translocating these signals from locally infected leaf to the uninfected parts of the plant. Consequently, *Trichoderma* helps in the accumulation of PR proteins and SA in the uninfected leaves, thus allowing the low nitrate stress plants to show enhanced resistance.

## Material and Methods

### Plant material and growth conditions

Seeds of *Arabidopsis thaliana* ecotype Columbia 0 (*Col0;* WT) were sown in plastic pots (6.5 × 7.5 cm) containing autoclaved soilrite: agropeat (1:1) mix and stratified at 4°C in the dark for 48 h. The pots were then kept in a growth room under short day conditions (8h-light, 16h-dark), 22/18°C (day/night) temperatures, relative humidity of 60% and 180-200 μE m^−2^ s^−1^ light intensity. Initially, plants were bottom irrigated for a week, once with half strength Hoagland’s solution and once with water. Then, 0.1 and 3 mM NO_3_^−^ concentrations were provided to the growing plants weekly. The NO_3_^−^ nutrient solution in Hoagland’s media contained either 0.1 mM or 3 mM KNO_3_, according to modified Hoagland’s nutrient solution (Hoagland and Arnon, 1950). 28-30 day-old plants with fully developed rosettes were used for the experiments. The seeds of *nrt2.1* (SALK_035429C) and *nia1,2* (N6936) were procured from ABRC.

### *Trichoderma* supplementation

*T. asperelloides* (*T203* strain) was grown on potato dextrose agar (PDA; Himedia) plates for 15 days under low light conditions until sporulation. Conidia were harvested by gently scraping the petridish and pouring 10 ml of sterile water over the surface and collecting the resultant suspension. Spores were evaluated up to 1□10^9^ spores/ml. It was thoroughly mixed into the soilrite mix and distributed into the individual plastic pots. WT plants in *Trichoderma* supplemented pots were also grown similarly as described above.

### Pathogen preparation and infiltration

*Pseudomonas syringae* pv. tomato DC3000 (*Pst*DC3000; *avrRpm1*, avirulent) were grown in King’s B (KB) medium containing 50 μg ml^−1^ rifampicin. Primary and secondary culture was prepared according to standard protocols (Liu *et al*. 2015). The avirulent bacterial density was adjusted to 2×10^7^ CFU ml^−1^ for LAR assay and primary inoculations and virulent *P. syringae* was used at 2×10^6^ CFU ml^−1^ for challenge inoculations. Mock infiltration (control) was performed with 10 mM MgCl_2_. *Pst* inoculations were made by syringe infiltration on the abaxial side of the leaves manually (Liu et al. 2015).

### LAR Assay

The procedure of the LAR assay is described schematically in Fig. **S3 a&b**. 30-day-old plants were used for this experimental set up and were transferred to an infection room prior to pathogen infiltration. Two sets of experiments were performed simultaneously, the first set were mock infiltrated (10 mM MgCl_2_) plants and the second set were the pathogen infiltrated (avirulent *Pst*DC3000; *avr*Rpm1) plants. The pathogen inoculated leaves were harvested at 0, 6, 24 and 48 hours post inoculation for RNA extraction and other assays. This assay was repeated more than three times in order to ensure the reproducibility of the results.

### SAR Assay

The procedure of the SAR assay is described schematically in Fig. **S3c**. The SAR assay was performed according to the protocol described in Cameron *et al*. (1999) in a specific infection room with appropriate growth conditions. Primary inoculation was performed with avirulent *Pst*DC3000 (*avrRpm1*) on one leaf per plant (marked with pink sticker; Fig. **S1b**) and the entire experimental set up was left for two days. Thereafter, a secondary (challenge) inoculation was performed with virulent *P. syringae* on four outer leaves (marked with black marker pen) other than primary inoculated leaf (Fig. **S1b**). Further, there were 5-6 non-inoculated distal leaves left per plant. Hereafter, the inoculated leaves are described as *Pst* or inoculated leaves and non-inoculated leaves are described as distal throughout the text. The inoculated and distal leaves from each treatment were harvested (Fig. **S3**) for RNA extraction and other experiments post challenge inoculation. This assay was repeated more than three times in order to obtain reproducible results.

### Electrolyte leakage

Leaf discs (5 mm diameter) were taken and electrolyte leakage was monitored exactly as described in (Gupta *et al*. 2013).

### *In planta* bacterial number quantification assay

The bacterial number in leaves from *Pst* treated plants during LAR and SAR responses were assessed was calculated as per Gupta *et al*. (2016).

### Determination of nitrate levels and nitrate uptake assays

Nitrate levels in leaves were determined using the protocol described in Hachiya and Okamoto (2017). The schematic representation of the experimental design and nitrate uptake assay in leaves is shown in Fig. **S4a and b**. Nitrate levels were also determined in 1 g each of S: A mixture, 0.1 mM N fed S: A mixture and 3 mM N fed S: A mixture (with and without *Trichoderma*; Fig. **S4c**). The procedure for nitrate determination in S: A mix and leaves is mentioned below.

#### Nitrate determination in S: A mix

For determining the basal nitrate levels, the individual pots were filled with S: A mixture (with and without *Trichoderma*) and 0.1 mM and 3 mM nitrate was supplied, allowed to soak for 2 hours and excess was drained off by pressing. The detailed protocol is described in Fig. **S4a and b** (Supplementary Information). Apparent nitrate was calculated from a standard curve of nitrate concentration (Fig. **S4d**) using the absorbance values at 410 nm.

#### Nitrate determination in leaves

Nitrate uptake was measured in leaves of 0.1 mM and 3 mM nitrate grown WT in the presence and absence of *Trichoderma* plants till 15 days and is represented as percent (%) nitrate uptake up to Day 15 (Fig. **S4b, Fig. 3a**). For each nitrate treatment, six pots containing healthy rosettes (22 d old) were filled with autoclaved S: A mixture (with and without *Trichoderma*). The detailed protocol is described in Supplementary Information of Fig. **S4b**.

### Expression profiling by qRT-PCR

Inoculated and non-inoculated leaves were immediately frozen in liquid nitrogen and stored at −80°C. RNA extraction, cDNA synthesis and qPCR was performed according to Wany *et al*. (2017; 2018). The synthesized cDNAs were used as templates in qRT-PCRs using the primers listed in Table S1. Fold change in the expression of the target genes was normalized to the *Arabidopsis* reference genes; ubiquitin, *18sRNA* (GQ380689) and *YSL8* (X69885.1). Fold expression relative to control treatment was determined by ΔΔCT values. Three biological experiments (with three independent replicates for each experiment) were performed for each treatment.

### NO estimation

For this experiment, NO was measured from the following five different combinations in roots; 1. WT; 2. WT + *T203 (Trichoderma);* 3. *nia1,2* double mutants + *T203*; 4. WT + cPTIO (carboxy-PTIO potassium salt); 5. WT + cPTIO + *T203*. WT and *nia1,2* plants were grown for one week in plates containing 0.1 mM and 3 mM NO_3_^−^ concentrations. A spore suspension of *Trichoderma* was poured over these 7d old plants and incubated for 2 minutes, 10 minutes and 24 hours, respectively. Then, the roots were incubated in 10 μM DAF-FM DA (4-amino-5-methylamino-2’,7’-difluorofluorescein diacetate) in 100 mM HEPES buffer (pH 7.2) placed in a 1.5 ml tube, incubated for 15 minutes in the dark and photographed using a fluorescence microscope (Nikon*80i*, Japan) set at 495 nm excitation and 515 nm emission wavelengths.

### Determination of ROS levels

Production of hydrogen peroxide (H_2_O_2_) in inoculated and distal leaves was detected by Diaminobenzidine tetrahydrochloride (DAB) staining as per Daudi *et al*. (2012) Superoxide levels were measured by *in vivo* staining with Nitroblue tetrazolium chloride (NBT, SA, USA) (Jambunathan, 2010).

### Histochemical detection of HR

Hypersensitive cell death in inoculated leaves was visualized by the trypan blue staining method according to Fernández-Bautista *et al*. (2002).

### SA levels

The SA levels were measured by HPLC according to the protocol described in Singh *et al*. (2013).

## Acknowledgements

We thank Dr. Yariv Brotman for providing the *T203* strain. Seeds of *nahG* and *npr1* mutants were provided by Prof. Ashis Nandi, JNU. Seeds of *nsHb^+^ over-expressing transgenic lines* was provided by Dr. Kim Hebelstrup. This research was funded by SERB, DST (NPDF to AW), UGC (SRF to PKP) and SERB-ECR and DBT-IYBA award to KJG. CIF of NIPGR is greatly acknowledged.

## Supplementary/Supporting Information

### Figure Legends of Supplementary files

**Fig. S1**: Phenotype of plants grown in different nitrate nutrition pre- and post-challenge inoculation **(a)** The growth pattern and phenotype of WT and *Trichoderma* treated WT plants in 3 mM and 0.1 mM NO_3_^−^ nutrition; **(b)** Effect of post-secondary challenge on plant phenotype, 1°-primary inoculation, 2°-secondary challenge and D-Distal leaves; **(c)** Hyponastic response (petiole elongation) shown by 0.1 mM and 3 mM NO_3_^−^ plants after primary inoculation.

**Fig. S2:** Electrolyte leakage of mock plants and morphological growth parameters during SAR **(a)** Electrolyte leakage from mock infiltrated plants in SAR; Morphological growth parameters observed in WT and treated WT plants; **(b)** Leaf number; **(c)** Biomass measured as fresh weight (FW); **(d)** Total chlorophyll content in *Pst* treated and untreated plants.

**Fig. S3:** Experimental design of LAR and SAR assay

**(a)** Description of plant samples taken for study and description of plants in absolute control, control and treatment. The sample size for each pot described here is n= 20-25 and plant’s age was 30 days old.

**(b)** In LAR assay, there were two sets of experiments performed at the same time. Set 1 comprised of mock infiltrated plants, most of the healthy leaves of the plants are syringe infiltrated by 10 mM MgCl_2_. Set 2 comprised of pathogen infiltrated plants, most of the healthy leaves were syringe infiltrated by avirulent *Pst*DC3000. The plants were left to acclimatize for 1 hour and then sample collection was performed at 0, 6, 24 and 48 hours post inoculation (hpi). Lowermost bar also shows the age of the plant during the experiment. All the infections were performed in infection room under standard conditions.

(c) In SAR assay, there were two sets of experiments performed at the same time. Set 1 comprised of plants initially infiltrated with primary inoculation of avirulent (*Pst*DC3000;*avr*Rpm1) and then secondary or challenge inoculation with virulent *Pst*DC3000. Set 2 comprised of mock infiltrated plants with both primary and secondary inoculation with 10 mM MgCl_2_. The incubation time for each inoculation is described in the figure. The time of primary and secondary inoculation was same (i.e. 10 am) and time selected for sampling is after the completion of incubation of challenge inoculation. The samples were collected after completing 3 days of challenge inoculation are denoted as 0 (10 am), 6 (4 pm), 24 (10 am next day) and 48 (10 am next to next day) hours post challenge (hpc).

Fig. S4 Experimental design and methodology of nitrate determination.

(a) Schematic representation of nitrate determination experimental design is shown here. First, the nitrate levels were determined in absolute control and soilrite: agropeat mix (with and without *Trichoderma*) according to Hachiya and Okamoto, 2017. 1 g of these S: A mixes were taken in 50 ml centrifuge tubes and immediately added pre-heated 10 ml of ultrapure water. Heat will denature nitrate reductase present in S: A mix and will be unable to utilize available nitrate in the substrate. Gently vortexed the tubes and kept at 100°C water bath and shaken every 5 minutes. After incubation, allowed the samples to cool down and settle down completely. Decanted the supernatant in another fresh tube. Centrifuged at maximum speed for 20 minutes at RT. Gently removed the clear supernatant and measured nitrate according to assay described in Hachiya and Okamoto (2017).

(b) Nitrate uptake assay was evaluated in rosette leaves of WT and WT+T plants (with and without *Trichoderma*) in a day-wise set up and percent nitrate uptake up to Day 15 is calculated. The trays containing pots were bottom irrigated with 0.1 mM and 3 mM nitrate nutrient Hoagland’s solution, allowed the pots to soak the entire nutrient solution for 2 hours and excess nutrient solution was drained off by pressing the S: A mixture. The leaves were sampled from Day 0 till Day 15 (even days) for nitrate uptake assay in the same set of plants (one leaf from each pot each day of sampling). During this period, nutrient was not supplied to the plants, they were only watered to ensure no drying of the S: A mixture and cause no drought to plants. A separate set of 6 pots of WT rosettes (absolute control) was also kept for calibrating the basal levels of nitrate present in leaves without flooding with 0.1 or 3 mM nitrate solution. The nitrate levels (leaves) obtained from this absolute control were subtracted from the nitrate treated pots. Each day-wise nitrate levels were calculated by subtracting the values of Day 2, 4, 6, 8, 10, 12 and 15 from Day 0 nitrate levels and subsequently % nitrate uptake was evaluated, shown in Fig. 3(a).

(c) Nitrate levels were also determined in 1 g of soilrite: agropeat mixture (blue bar), 0.1 mM fed S: A mix (light orange bar), 3 mM fed S: A mix (light green bar), 0.1 mM fed S: A mix with *Trichoderma* (light purple bar) and 3 mM fed S: A mix with *Trichoderma* (yellow bar). Data are average mean values ± SE with n=3. Statistical significance was tested by one-way ANOVA followed by Dunnett’s multiple comparison test. The different letters above each column represent significance difference between means at p□<□0.05.

(d) Standard curve of potassium nitrate used to calculate apparent nitrate concentration. Nitrate standard series of 2, 4, 6, 8, 10 and 12 mM were prepared and assay was performed according to Hachiya and Okamoto, (2017). A straight line curve generated was used to determine the nitrate concentration (mM) using the formula = Abs/0.055, where 0.055 is the value according to straight line curve.

**Fig S5**: **(a)** Electrolyte leakage in distal leaves in WT and WT+T leaves during SAR **(b)** % cell death quantified in Trypan blue images of both inoculated (Pst) and uninoculated (distal) leaves in WT and WT+T leaves during SAR using ImageJ. Results are presented as % necrotized leaf area compared with the total surface of leaf analyzed by ImageJ (ver 3.2) and represent means ± SE of 6 leaves per treatment.

**Table S1:**
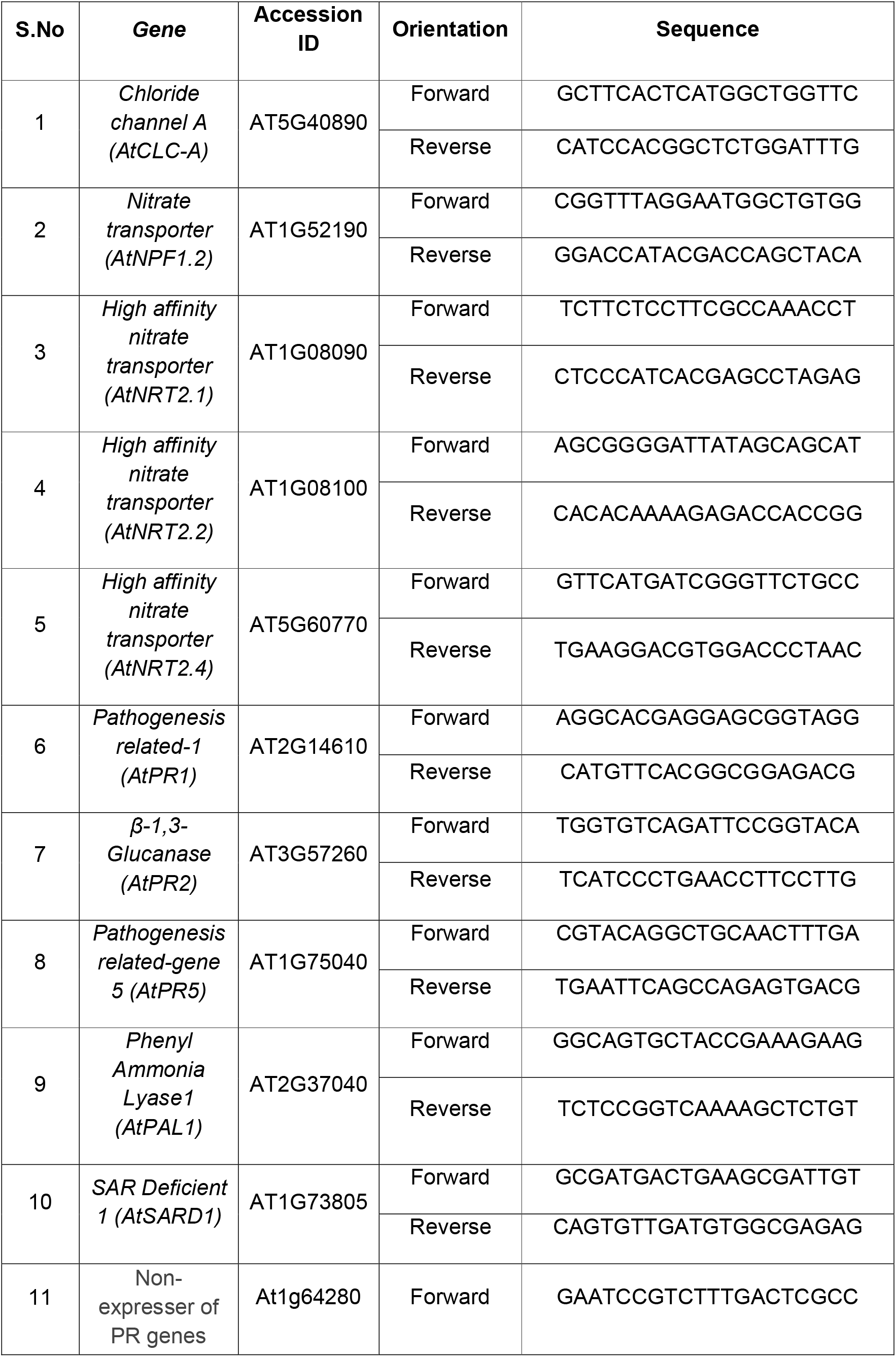
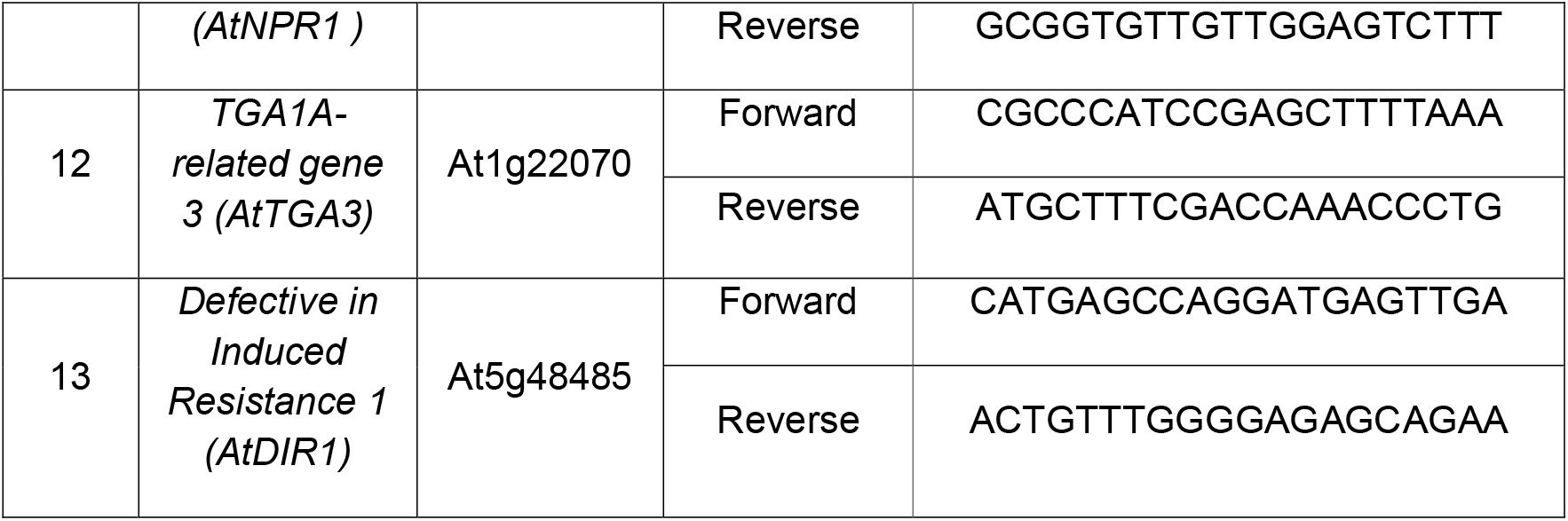
List of primers

## Notes

### Competing Interest Statement

The authors have declared no competing interest.

### Summary of Updates

Nitrogen (N) is essential for growth, development and defense but, how low N affects defense and the role of Trichoderma in enhancing defense under low nitrate is not known. Low nitrate fed Arabidopsis plants displayed reduced growth and compromised local and systemic acquired resistance responses when infected with both avirulent and virulent Pseudomonas syringae DC3000. These responses were enhanced in the presence of Trichoderma. The mechanism of increased local and systemic acquired resistance mediated by Trichoderma involved increased N uptake and enhanced protein levels via modulation of nitrate transporter genes. The nrt2.1 mutant is compromised in local and systemic acquired resistance responses suggesting a link between enhanced N transport and defense. Enhanced N uptake was mediated by Trichoderma elicited nitric oxide (NO). Low NO producing nia1,2 mutant and nsHb+ over expressing lines were unable to induce nitrate transporters and thereby compromised defense in the presence of Trichoderma under low N suggesting a signaling role of Trichoderma elicited NO. Trichoderma also induced SA and defense gene expression under low N. The SA deficient NahG transgenic line and the npr1 mutant were also compromised in Trichoderma-mediated local and systemic acquired resistance responses. Collectively our results indicated that the mechanism of enhanced plant defense under low N mediated by Trichoderma involves NO, ROS, SA production as well as the induction of NRT and marker genes for systemic acquired resistance.

## References

Astier, J., Gross, I. and Durner, J. (2018) Nitric oxide production in plants: an update. J. Exp. Bot. 69, 3401–3411.

Alvarez, M. E., Pennell, R. I., Meijer, P. J. et al. (1998) Reactive oxygen intermediates mediate a systemic signal network in the establishment of plant immunity. Cell. 92, 773–784.

Alves, M., Dadalto, S., Gonçalves, A. et al. (2014) Transcription factor functional protein-protein interactions in plant defense responses. Proteomes. 2, 85–106.

Asmawati, L., Widiastuti, A. and Sumardiyono, C. (2017) Induction of Reactive Oxygen Species by *Trichoderma* spp. Against Downy Mildew in Maize. In Proceedings of the 1st International Conference on Tropical Agriculture. Springer, Cham:139–146.

Boudsocq, S., Niboyet, A. and Lata, J. C. et al. (2012) Plant preference for ammonium versus nitrate: a neglected determinant of ecosystem functioning?. Am Nat. 180, 60–69.

Brotman, Y., Lisec, J. and Méret, M. et al. (2012) Transcript and metabolite analysis of the Trichoderma-induced systemic resistance response to Pseudomonas syringae in *Arabidopsis thaliana*. Microbiology. 158, 139–146.

Brotman, Y., Gupta, K. J. and Vetribo, A. (2010) *Trichoderma*: Quick Guide. Curr Biol. 20, 390–391.

Calderon, A. A., Zapata, J. M. and Barcelo, A. R. (1994) Peroxidase-mediated formation of resveratrol oxidation products during the hypersensitive-like reaction of grapevine cells to an elicitor from Trichoderma viride. Physiol Mol Plant Path. 44, 289–299.

Cameron, R. K., Dixon, R. A. and Lamb, C. J. (1994) Biologically induced systemic acquired resistance in Arabidopsis thaliana. Plant J. 5, 715–725.

Cameron, R. K., Paiva, N. L., Lamb, C. J. et al. (1999) Accumulation of salicylic acid and *PR-1* gene transcripts in relation to the systemic acquired resistance (SAR) response induced by *Pseudomonas syringae* pv. tomato in Arabidopsis. Physiol Mol Plant Path. 55, 121–130.

Cao, H., Glazebrook, J., Clarke, J. D. et al. (1997) The Arabidopsis NPR1 gene that controls systemic acquired resistance encodes a novel protein containing ankyrin repeats. Cell. 88, 57–63.

Carella, P., Isaacs, M. and Cameron, R. K. (2015) Plasmodesmata-located protein overexpression negatively impacts the manifestation of systemic acquired resistance and the long-distance movement of Defective in Induced Resistance1 in A rabidopsis. Plant Biol. 17, 395–401.

Champigny, M. J., Isaacs, M., Carella, P. et al. (2013) Long distance movement of DIR1 and investigation of the role of DIR1-like during systemic acquired resistance in Arabidopsis. Front Plant Sci. 4, 230.

Contreras-Cornejo, H. A., Macías-Rodríguez, L., del-Val, E. et al. (2016) Ecological functions of Trichoderma spp. and their secondary metabolites in the rhizosphere: interactions with plants. FEMS Microbiol Ecol. 92, fiw036.

Daudi, A. and O’Brien, J. A. 2012. Detection of hydrogen peroxide by DAB staining in Arabidopsis leaves. Bio-Protocol. 2, 1–4.

Dechorgnat, J., Patrit, O., Krapp, A. et al. (2012) Characterization of the Nrt2. 6 gene in *Arabidopsis thaliana*: a link with plant response to biotic and abiotic stress. PloS one. 7, e42491.

Delaney, T. P., Uknes, S., Vernooij, B. et al. (1994) A central role of salicylic acid in plant disease resistance. Science. 266, 1247–1250.

Delledonne, M., Xia, Y., Dixon, R. A. et al. (1998) Nitric oxide functions as a signal in plant disease resistance. Nature. 394, 585.

Dietrich, R., K. Ploss, and M. Heil. (2004) Constitutive and induced resistance to pathogens in *Arabidopsis thaliana* depends on nitrogen supply. Plant Cell Environ. 27, 896–906.

Domínguez, S., Rubio, M. B., Cardoza, R. E. et al. (2016) Nitrogen metabolism and growth enhancement in tomato plants challenged with *Trichoderma* harzianum expressing the *Aspergillus nidulans* acetamidase *amdS* gene. Front Microbiol. 7, 1182.

Dorey, S., Baillieul, F., Pierrel, M. A. et al. (1997) Spatial and temporal induction of cell death, defense genes, and accumulation of salicylic acid in tobacco leaves reacting hypersensitively to a fungal glycoprotein elicitor. Mol Plant Microbe. 10, 646–655.

Dorey, S., Kopp, M., Geoffroy, P. et al. (1999) Hydrogen peroxide from the oxidative burst is neither necessary nor sufficient for hypersensitive cell death induction, phenylalanine ammonia lyase stimulation, salicylic acid accumulation, or scopoletin consumption in cultured tobacco cells treated with elicitin. Plant Physiol. 121, 163–172.

Durner, J. and Klessig, D. F. (1999) Nitric oxide as a signal in plants. Curr Opin Plant Biol. 2, 369–374.

Fagard, M., Launay, A., Clément, G. et al. (2014) Nitrogen metabolism meets phytopathology. J Exp Bot. 65, 5643–5656.

Fernández-Bautista, N., Domínguez-Núñez, J. A., Moreno, M. C. et al. (2002) Plant tissue trypan blue staining during phytopathogen infection. Bio-Protocol. 6, e2078.

Frungillo, L., de Oliveira, J. F. P., Saviani, E. E. et al. (2013) Modulation of mitochondrial activity by *S-nitrosoglutathione reductase* in *Arabidopsis thaliana* transgenic cell lines. BBA-Bioenergetics. 1827, 239–247.

Garnica-Vergara, A., Barrera-Ortiz, S., Muñoz-Parra, E. et al. (2016) The volatile 6-pentyl-2H-pyran-2-one from Trichoderma atroviride regulates Arabidopsis thaliana root morphogenesis via auxin signaling and ETHYLENE INSENSITIVE 2 functioning. New Phytol. 209, 1496–1512.

Gupta, A., Dixit, S. K. and Senthil-Kumar, M. (2016) Drought stress predominantly endures Arabidopsis thaliana to *Pseudomonas syringae* infection. Front. Plant Sci. 7, 808.

Gupta, K. J., Mur, L. A. and Brotman, Y. (2014) *Trichoderma asperelloides* suppresses nitric oxide generation elicited by *Fusarium oxysporum* in Arabidopsis roots. Mol Plant Microbe Interact. 27, 307–314.

Gupta, K. J., Brotman, Y., Segu, S. et al. (2013) The form of nitrogen nutrition affects resistance against *Pseudomonas syringae pv. phaseolicola* in tobacco. J Exp Bot. 64, 553–568.

Hachiya, T. and Okamoto, Y. (2017) Simple spectroscopic determination of nitrate, nitrite, and ammonium in Arabidopsis thaliana. Bio-Protocol. 7.

Hoagland, D. R. and Arnon, D. I. (1950) The water-culture method for growing plants without soil. Circular. Calif. Agr. Exp. Sta. Cir. 347.

Jambunathan, N. (2010) Determination and detection of reactive oxygen species (ROS), lipid peroxidation, and electrolyte leakage in plants. Methods Mol Biol. 639, 292–298.

Jones, J. D. and Dangl, J. L. (2006) The plant immune system. Nature. 444, 323.

Johnson, N. D., Liu, B. and Bentley, B. L. (1987) The effects of nitrogen fixation, soil nitrate, and defoliation on the growth, alkaloids, and nitrogen levels of *Lupinus succulentus* (Fabaceae). Oecologia. 74, 425–431.

Kim, D. S. and Hwang, B. K. (2014) An important role of the pepper phenylalanine ammonia-lyase gene (PAL1) in salicylic acid-dependent signalling of the defence response to microbial pathogens. J Exp Bot. 65, 2295–2306.

Krapp, A., David, L. C., Chardin, C. et al. (2014) Nitrate transport and signalling in Arabidopsis. J Exp Bot. 65, 789–798.

Lamb, C. and Dixon, R. A. (1997) The oxidative burst in plant disease resistance. Annu Rev Plant Biol. 48, 251–275.

Landrein, B., Formosa-Jordan, P., Malivert, A. et al. (2018) Nitrate modulates stem cell dynamics in Arabidopsis shoot meristems through cytokinins. PNAS USA. 115, 1382–1387.

Li, H., Hu, B. and Chu, C. (2017) Nitrogen use efficiency in crops: lessons from Arabidopsis and rice. J Exp Bot. 68, 2477–2488.

Liu, Y. and von Wirén, N. (2017) Ammonium as a signal for physiological and morphological responses in plants. J Exp Bot. 68, 2581–2592.

Maldonado, A. M., Doerner, P., Dixon, R. A. et al. (2002) A putative lipid transfer protein involved in systemic resistance signalling in Arabidopsis. Nature. 419, 399–403.

Martínez-Medina, A., Fernández, I., Sánchez-Guzmán, M. J. et al. (2013) Deciphering the hormonal signalling network behind the systemic resistance induced by Trichoderma harzianum in tomato. Front Plant Sci. 4, 206.

Métraux, J. P., Signer, H., Ryals, J. et al. (1990) Increase in salicylic acid at the onset of systemic acquired resistance in cucumber. Science, 250, 1004–1006.

Mur, L. A., Brown, I. R., Darby, R. M. et al. (2000) A loss of resistance to avirulent bacterial pathogens in tobacco is associated with the attenuation of a salicylic acid-potentiated oxidative burst. Plant J. 23, 609–621.

Mur, L. A., Kenton, P., Lloyd, A. J. et al. (2008) The hypersensitive response; the centenary is upon us but how much do we know?. J Exp Bot. 59, 501–520.

Mur, L. A., Simpson, C., Kumari, A. et al. (2017) Moving nitrogen to the centre of plant defence against pathogens. Ann. Bot. 119, 703–709.

O’Brien, J. A., Vega, A., Bouguyon, E. et al. (2016) Nitrate transport, sensing, and responses in plants. Mol Plant. 9, 837–856.

Oliva, R. and Quibod, I. L. (2017) Immunity and starvation: new opportunities to elevate disease resistance in crops. Curr Opin Plant Biol. 38, 84–91.

Park, S. W., Kaimoyo, E., Kumar, D. et al. (2007). Methyl salicylate is a critical mobile signal for plant systemic acquired resistance. Science. 318, 113–116.

Pieterse, C. M., Van Wees, S. C., Hoffland, E. et al. (1996) Systemic resistance in Arabidopsis induced by biocontrol bacteria is independent of salicylic acid accumulation and pathogenesis-related gene expression. Plant Cell. 8, 1225–1237.

Pieterse, C. M., Zamioudis, C., Berendsen, R. L. et al. (2014) Induced systemic resistance by beneficial microbes. Annu Rev Phytopathol. 52, 347–375.

Ryals, J. A., Neuenschwander, U. H., Willits, M. G. et al. (1996) Systemic acquired resistance. Plant cell. 8, 1809.

Shoresh, M., Harman, G. E. and Mastouri, F. (2010) Induced systemic resistance and plant responses to fungal biocontrol agents. Annu Rev Phytopathol. 48, 21–43.

Shoresh, M., Yedidia, I. and Chet, I. (2005) Involvement of jasmonic acid/ethylene signaling pathway in the systemic resistance induced in cucumber by Trichoderma asperellum T203. Phytopathology. 95, 76–84.

Singh, V., Roy, S., Giri, M. K., Chaturvedi, R. et al. (2013) Arabidopsis thaliana FLOWERING LOCUS D is required for systemic acquired resistance. MPMI. 26, 1079–1088.

Snoeijers, S. S., Pérez-García, A., Joosten, M. H. et al. (2000) The effect of nitrogen on disease development and gene expression in bacterial and fungal plant pathogens. Eur J Plant Pathol. 106, 493–506.

Stout, M. J., Brovont, R. A. and Duffey, S. S. (1998) Effect of nitrogen availability on expression of constitutive and inducible chemical defenses in tomato, *Lycopersicon esculentum*. J. Chem. Ecol. 24, 945–963.

Torres, M. A., Dangl, J. L. and Jones, J. D. (2002) Arabidopsis gp91phox homologues AtrbohD and AtrbohF are required for accumulation of reactive oxygen intermediates in the plant defense response. PNAS. 99, 517–522.

Torres, M. A., Jones, J. D. and Dangl, J. L. (2005) Pathogen-induced, NADPH oxidase–derived reactive oxygen intermediates suppress spread of cell death in Arabidopsis thaliana. Nat. Genet. 37, 1130–1134.

Tsay, Y. F., Chiu, C. C., Tsai, C. B. et al. (2007) Nitrate transporters and peptide transporters. FEBS Lett. 581, 2290–2300.

Walker, R. L., Burns, I. G. and Moorby, J. (2001) Responses of plant growth rate to nitrogen supply: a comparison of relative addition and N interruption treatments. J Exp Bot. 52, 309–317.

Ward, J. L., Forcat, S., Beckmann, M. et al. (2010) The metabolic transition during disease following infection of *Arabidopsis thaliana* by *Pseudomonas syringae pv. tomato*. Plant J. 63, 443–4

Wang, L., Tsuda, K., Truman, W. et al. (2011) CBP60g and SARD1 play partially redundant critical roles in salicylic acid signaling. Plant J. 67, 1029–1041.

Wany, A., Gupta, A. K., Kumari, A. et al. (2019) Nitrate nutrition influences multiple factors in order to increase energy efficiency under hypoxia in Arabidopsis. Ann Bot. 123, 691–705.

Wany, A., Kumari, A., and Gupta, K. J. (2017) Nitric oxide is essential for the development of aerenchyma in wheat roots under hypoxic stress. Plant, cell environ. 40, 3002–3017.

Yedidia, I., Shoresh, M., Kerem, Z. et al. (2003) Concomitant induction of systemic resistance to Pseudomonas syringae pv. lachrymans in cucumber by Trichoderma asperellum (T-203) and accumulation of phytoalexins. Appl. Environ. Microbiol. 69, 7343–7353.

Zhang, Y., Xu, S., Ding, P. et al. (2010) Control of salicylic acid synthesis and systemic acquired resistance by two members of a plant-specific family of transcription factors. PNAS. 107, 18220–18225.

