## Supplementary material for "*Trichoderma asperelloides* enhances local and systemic acquired resistance response under low nitrate nutrition in Arabidopsis": Suppliementary data

Supporting Information

Article title: *Trichoderma asperelloides* enhances local (LAR) and systemic acquired resistance (SAR) response under low nitrate nutrition in Arabidopsis

Authors: Aakanksha Wany, Pradeep K. Pathak, Alisdair R Ferine Kapuganti Jagadis Gupta*

**Fig S1**: Phenotype of plants grown in different nitrate nutrition pre- and post-challenge inoculation **(a)** The growth pattern and phenotype of WT and Trichoderma treated WT plants in 3 mM and 0.1 mM NO_3_^-^ nutrition; **(b)** Effect of post-secondary challenge on plant phenotype, 1°- primary inoculation, 2°-secondary challenge and D-Distal leaves; **(c)** Hyponastic response (petiole elongation) shown by 0.1 mM and 3 mM NO_3_^-^ plants after primary inoculation.

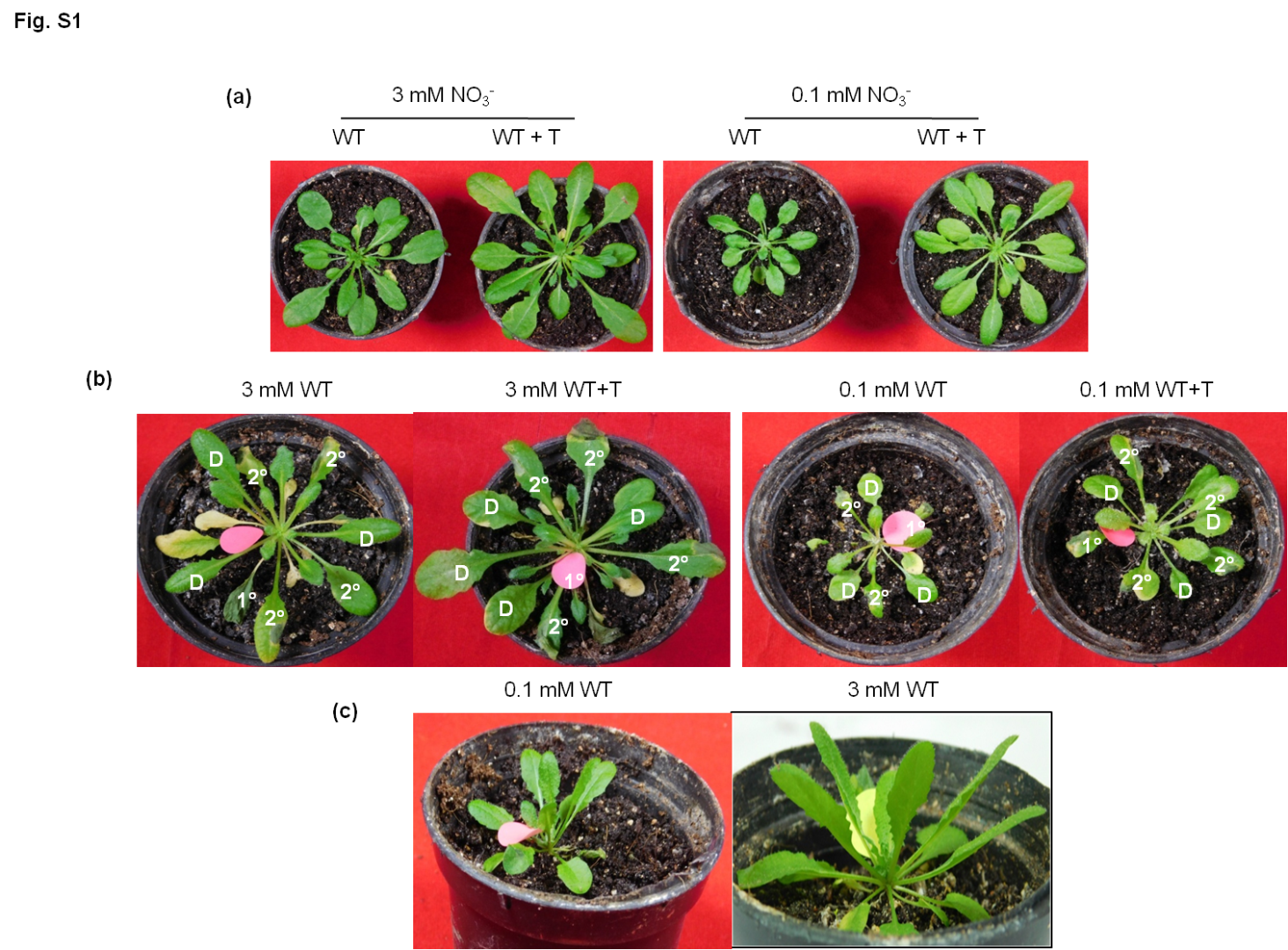

**Fig. S2:** Electrolyte leakage of mock plants and morphological growth parameters during SAR **(a)** Electrolyte leakage from mock infiltrated plants in SAR; Morphological growth parameters observed in WT and Trichoderma treated WT plants; **(b)** Leaf number; **(c)** Biomass measured as fresh weight (FW); **(d)** Total chlorophyll content in *Pst* treated and untreated plants.

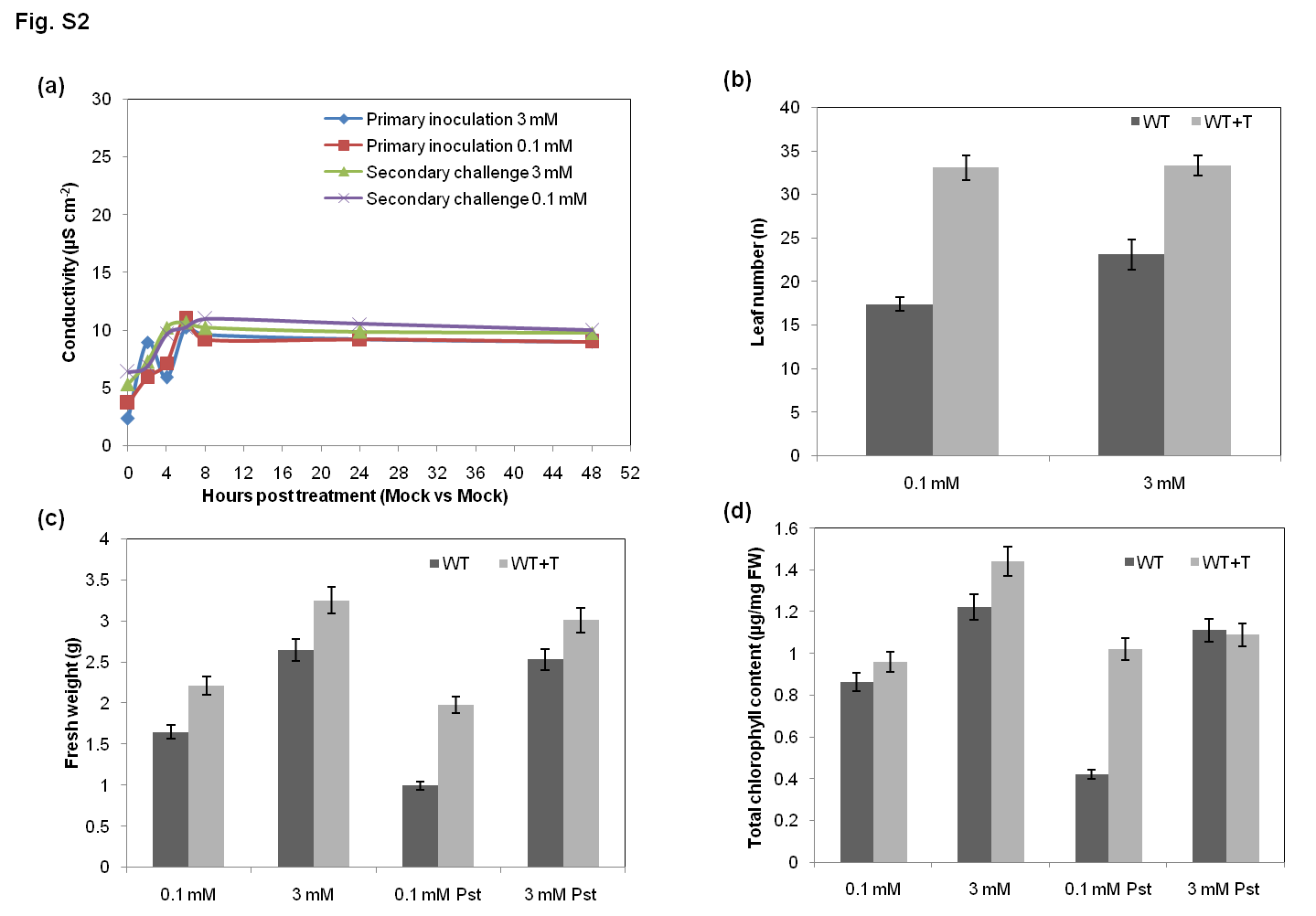

**Fig. S3: Experimental design of LAR and SAR assay**

**(a)** Description of plant samples taken for study and description of plants in absolute control, control and treatment. The sample size for each pot described here is n= 20-25 and plant's age was 30 days old.

**(b)** In LAR assay, there were two sets of experiments performed at the same time. Set 1 comprised of mock infiltrated plants, most of the healthy leaves of the plants are syringe infiltrated by 10 mM MgCl_2_. Set 2 comprised of pathogen infiltrated plants, most of the healthy leaves were syringe infiltrated by avirulent *Pst*DC3000. The plants were left to acclimatize for 1 hour and then sample collection was performed at 0, 6, 24 and 48 hours post inoculation (hpi). Lowermost bar also shows the age of the plant during the experiment. All the infections were performed in infection room under standard conditions.

(c) In SAR assay, there were two sets of experiments performed at the same time. Set 1 comprised of plants initially infiltrated with primary inoculation of avirulent (*Pst*DC3000;*avr*Rpm1) and then secondary or challenge inoculation with virulent *Pst*DC3000. Set 2 comprised of mock infiltrated plants with both primary and secondary inoculation with 10 mM MgCl_2_. The incubation time for each inoculation is described in the figure. The time of primary and secondary inoculation was same (i.e. 10 am) and time selected for sampling is after the completion of incubation of challenge inoculation. The samples were collected after completing 3 days of challenge inoculation are denoted as 0 (10 am), 6 (4 pm), 24 (10 am next day) and 48 (10 am next to next day) hours post challenge (hpc).

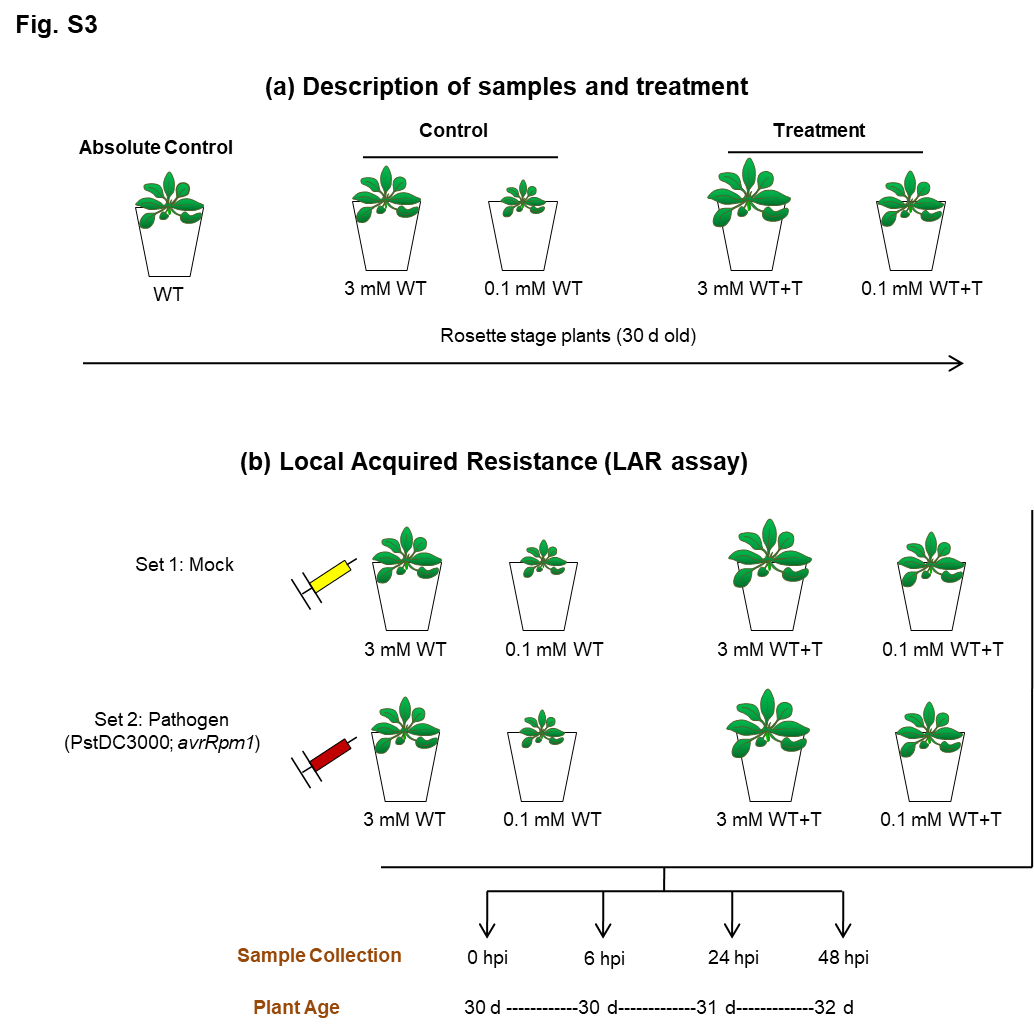

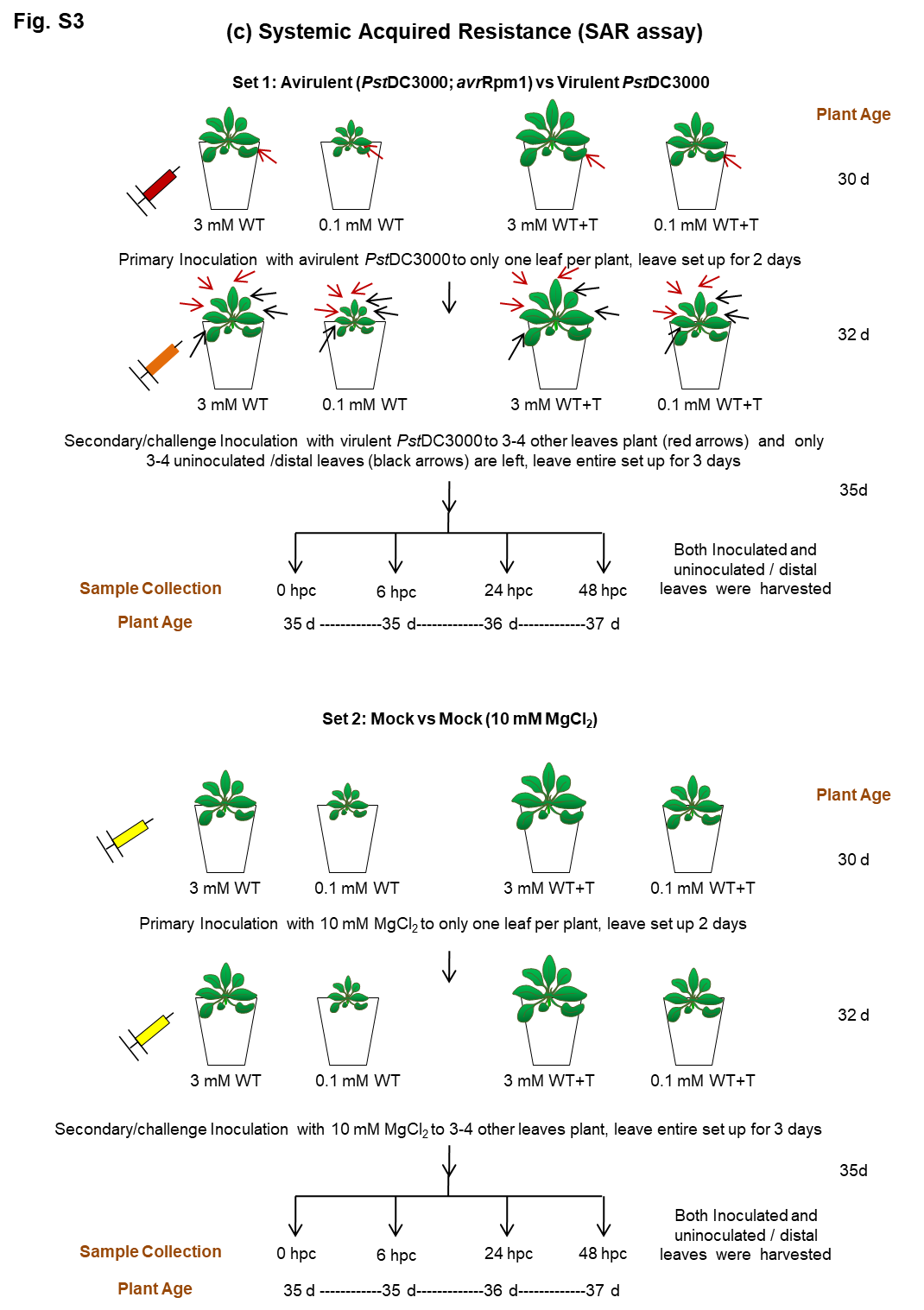

**Fig. S4** Experimental design and methodology of nitrate determination.

(a) Schematic representation of nitrate determination experimental design is shown here. First, the nitrate levels were determined in absolute control and soilrite:agropeat mix (with and without Trichoderma) according to Hachiya and Okamoto, 2017. 1 g of these S: A mixes were taken in 50 ml centrifuge tubes and immediately added pre-heated 10 ml of ultrapure water. Heat will denature nitrate reductase present in S: A mix and will be unable to utilize available nitrate in the substrate. Gently vortexed the tubes and kept at 100°C water bath and shaken every 5 minutes. After incubation, allowed the samples to cool down and settle down completely. Decanted the supernatant in another fresh tube. Centrifuged at maximum speed for 20 minutes at RT. Gently removed the clear supernatant and measured nitrate according to assay described in Hachiya and Okamoto (2017).
(b) Nitrate uptake assay was evaluated in rosette leaves of WT and WT+T plants (with and without Trichoderma) in a day-wise set up and percent nitrate uptake upto Day 15 is calculated. The trays containing pots were bottom irrigated with 0.1 mM and 3 mM nitrate nutrient Hoagland's solution, allowed the pots to soak the entire nutrient solution for 2 hours and excess nutrient solution was drained off by pressing the S: A mixture. The leaves were sampled from Day 0 till Day 15 (even days) for nitrate uptake assay in the same set of plants (one leaf from each pot each day of sampling). During this period, nutrient was not supplied to the plants, they were only watered to ensure no drying of the S: A mixture and cause no drought to plants. A separate set of 6 pots of WT rosettes (absolute control) was also kept for calibrating the basal levels of nitrate present in leaves without flooding with 0.1 or 3 mM nitrate solution. The nitrate levels (leaves) obtained from this absolute control were subtracted from the nitrate treated pots. Each day-wise nitrate levels were calculated by subtracting the values of Day 2, 4, 6, 8, 10, 12 and 15 from Day 0 nitrate levels and subsequently % nitrate uptake was evaluated, shown in Fig. 3(a).
(c) Nitrate levels were also determined in 1 g of soilrite: agropeat mixture (blue bar), 0.1 mM fed S:A mix (light orange bar), 3 mM fed S:A mix (light green bar), 0.1 mM fed S:A mix with Trichoderma (light purple bar) and 3 mM fed S:A mix with Trichoderma (yellow bar). Data are average mean values ± SE with n=3. Statistical significance was tested by one-way ANOVA followed by Dunnett's multiple comparison test. The different letters above each column represent significance difference between means at p < 0.05.
(d) Standard curve of potassium nitrate used to calculate apparent nitrate concentration. Nitrate standard series of 2, 4, 6, 8, 10 and 12 mM were prepared and assay was performed according to Hachiya and Okamoto, (2017). A straight line curve generated was used to determine the nitrate concentration (mM) using the formula = Abs/0.055, where 0.055 is the value according to straight line curve.

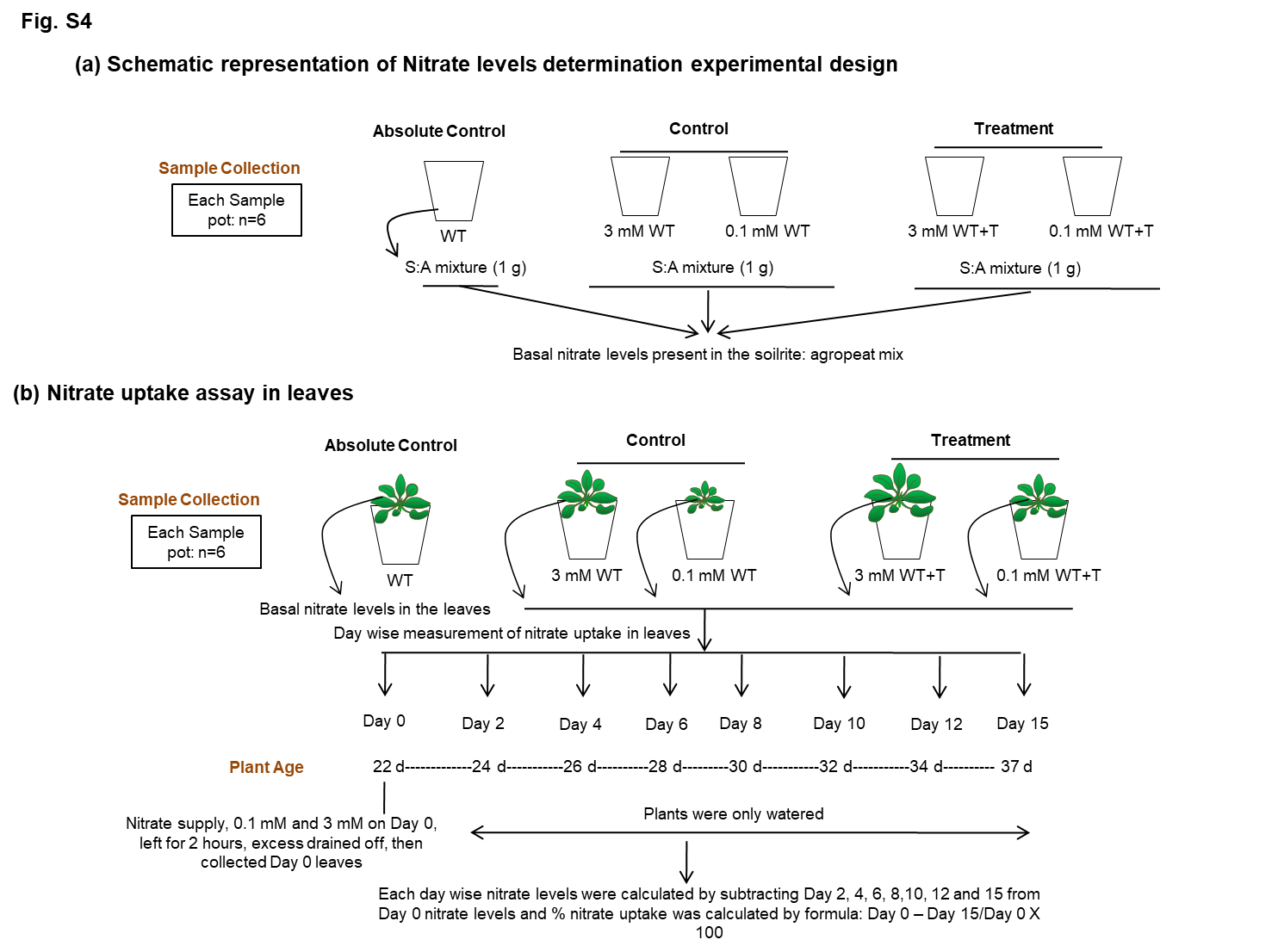

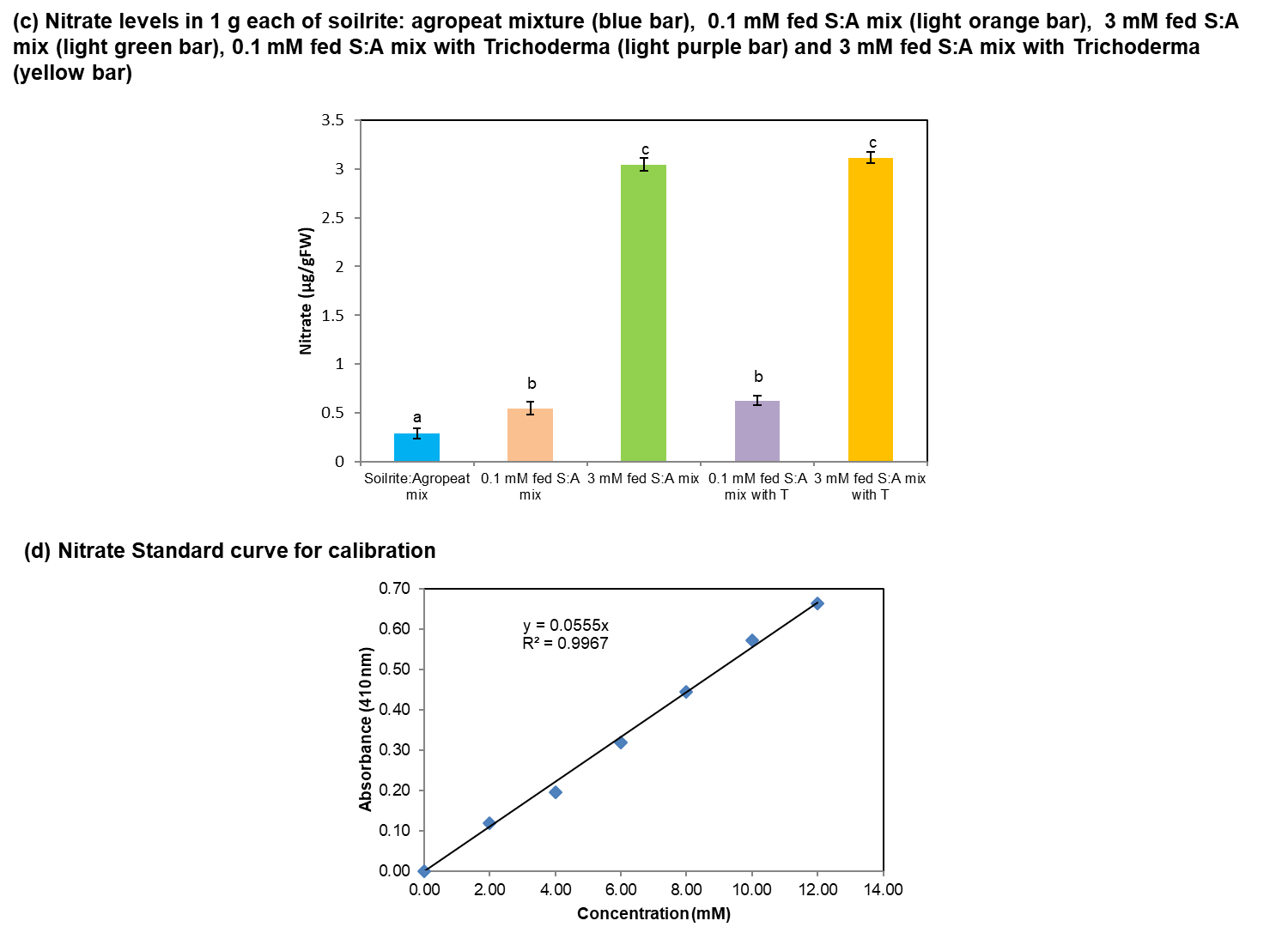

**Fig S5**: **(a)** Electrolyte leakage in distal leaves in WT and WT+T leaves during SAR **(b)** % cell death quantified in Trypan blue images of both inoculated (Pst) and uninoculated (distal) leaves in WT and WT+T leaves during SAR using ImageJ. Results are presented as % necrotized leaf area compared with the total surface of leaf analyzed by ImageJ (ver 3.2) and represent means ± SE of 6 leaves per treatment.

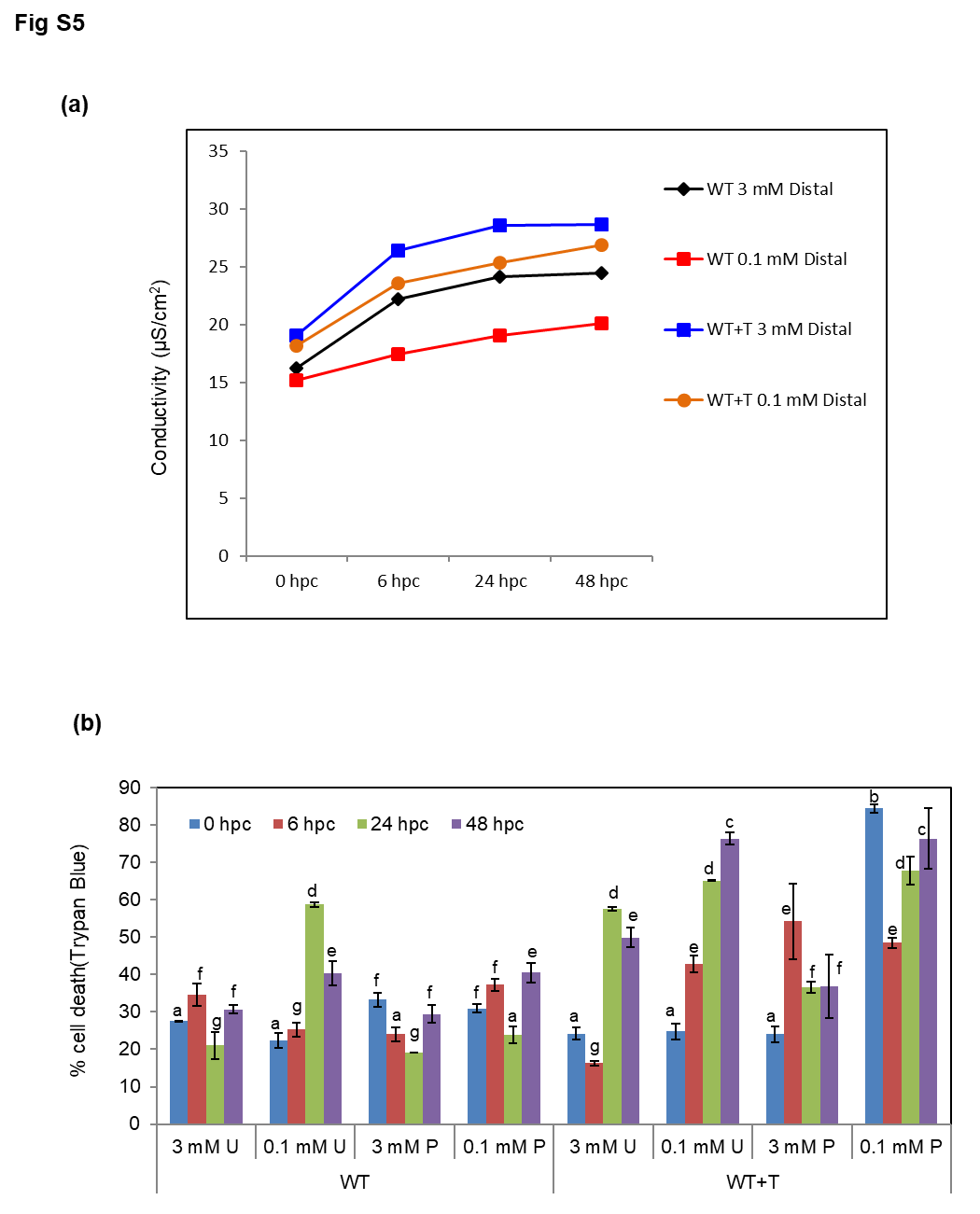

**Table S1**: List of primers

| **S.No** | ***Gene*** | **Accession ID** | **Orientation** | **Sequence** |
| --- | --- | --- | --- | --- |
| 1 | *Chloride channel A (AtCLC-A)* | AT5G40890 | Forward | GCTTCACTCATGGCTGGTTC |
|  |  |  | Reverse | CATCCACGGCTCTGGATTTG |
| 2 | *Nitrate transporter (AtNPF1.2)* | AT1G52190 | Forward | CGGTTTAGGAATGGCTGTGG |
|  |  |  | Reverse | GGACCATACGACCAGCTACA |
| 3 | *High affinity nitrate transporter (AtNRT2.1)* | AT1G08090 | Forward | TCTTCTCCTTCGCCAAACCT |
|  |  |  | Reverse | CTCCCATCACGAGCCTAGAG |
| 4 | *High affinity nitrate transporter (AtNRT2.2)* | AT1G08100 | Forward | AGCGGGGATTATAGCAGCAT |
|  |  |  | Reverse | CACACAAAAGAGACCACCGG |
| 5 | *High affinity nitrate transporter (AtNRT2.4)* | AT5G60770 | Forward | GTTCATGATCGGGTTCTGCC |
|  |  |  | Reverse | TGAAGGACGTGGACCCTAAC |
| 6 | *Pathogenesis related-1 (AtPR1)* | AT2G14610 | Forward | AGGCACGAGGAGCGGTAGG |
|  |  |  | Reverse | CATGTTCACGGCGGAGACG |
| 7 | *β-1,3-Glucanase (AtPR2)* | AT3G57260 | Forward | TGGTGTCAGATTCCGGTACA |
|  |  |  | Reverse | TCATCCCTGAACCTTCCTTG |
| 8 | *Pathogenesis related-gene 5 (AtPR5)* | AT1G75040 | Forward | CGTACAGGCTGCAACTTTGA |
|  |  |  | Reverse | TGAATTCAGCCAGAGTGACG |
| 9 | *Phenyl Ammonia Lyase1 (AtPAL1)* | AT2G37040 | Forward | GGCAGTGCTACCGAAAGAAG |
|  |  |  | Reverse | TCTCCGGTCAAAAGCTCTGT |
| 10 | *SAR Deficient 1 (AtSARD1)* | AT1G73805 | Forward | GCGATGACTGAAGCGATTGT |
|  |  |  | Reverse | CAGTGTTGATGTGGCGAGAG |
| 11 | Non-expresser of PR genes  *(AtNPR1 )* | At1g64280 | Forward | GAATCCGTCTTTGACTCGCC |
|  |  |  | Reverse | GCGGTGTTGTTGGAGTCTTT |
| 12 | *TGA1A-related gene 3 (AtTGA3)* | At1g22070 | Forward | CGCCCATCCGAGCTTTTAAA |
|  |  |  | Reverse | ATGCTTTCGACCAAACCCTG |
| 13 | *Defective in Induced Resistance 1 (AtDIR1)* | At5g48485 | Forward | CATGAGCCAGGATGAGTTGA |
|  |  |  | Reverse | ACTGTTTGGGGAGAGCAGAA |
